# Organization and control of the ascorbate biosynthesis pathway in plants

**DOI:** 10.1101/2020.06.23.167247

**Authors:** Mario Fenech, Vítor Amorim-Silva, Alicia Esteban del Valle, Dominique Arnaud, Araceli G. Castillo, Nicholas Smirnoff, Miguel A. Botella

**Affiliations:** Instituto de Hortofruticultura Subtropical y Mediterránea “La Mayora” (IHSM-UMA-CSIC), Departamento de Biología Molecular y Bioquímica, Facultad de Ciencias, Universidad de Málaga, Campus de Teatinos s/n, E-29071 Málaga, Spain; Biosciences, College of Life and Environmental Sciences, University of Exeter, Exeter, EX4 4QD, UK; Instituto de Hortofruticultura Subtropical y Mediterránea “La Mayora” (IHSM-UMA-CSIC), Departamento de Genética, Facultad de Ciencias, Universidad de Málaga, Campus de Teatinos s/n, E-29071 Málaga, Spain

**Author notes:** Author for Contact., Author for Contact Details: Miguel A. Botella, Instituto de Hortofruticultura Subtropical y Mediterránea “La Mayora” (IHSM-UMA-CSIC), Departamento de Biología Molecular y Bioquímica, Facultad de Ciencias, Universidad de Málaga, Campus de Teatinos s/n, E-29071 Málaga, Spain., Nicholas Smirnoff, Biosciences, College of Life and Environmental Sciences, University of Exeter, Exeter, EX4 4QD, UK. Mario Fenech, Vítor Amorim-Silva, Alicia Esteban del Valle, Dominique Arnaud, Araceli G. Castillo, Nicholas Smirnoff, Miguel A. Botella. Author contributions: M.F., V.A.S., M.A.B. and N.S. conceived the project and designed the experiments; M.F. performed the experiments and analyzed the data; N.S. performed the MCA simulation; V.A.S., M.A.B., and N.S. supervised the experiments; D.A. generated vtc2/VTC2-GFP transgenic lines and tested mutant complementation by AO assay; A.G.C. performed yeast two-hybrid assay with technical assistance by M.F.; A.E.V. provided technical assistance to M.F.; M.F., M.A.B. and N.S. wrote the article; V.A.S., M.A.B. and N.S. supervised and completed the writing; M.A.B. agrees to serve as the author responsible for contact and ensures communication. Dominique Arnaud current address: Laboratory of Molecular Plant Physiology and Functional Genomics and Proteomics of Plants, CEITEC-Central European Institute of Technology, Masaryk University, Kamenice 5, CZ-625 00, Brno, Czech Republic.

## Abstract

The enzymatic steps involved in l-ascorbate biosynthesis in photosynthetic organisms (the Smirnoff-Wheeler, SW pathway) has been well established and here we comprehensively analyze the subcellular localization, potential physical interactions of SW pathway enzymes and assess their role in control of ascorbate synthesis. Transient expression of GFP-fusions in *Nicotiana benthamiana* and Arabidopsis (*Arabidopsis thaliana*) mutants complemented with genomic constructs showed that while GME is cytosolic, VTC1, VTC2, VTC4, and l-GalDH have cytosolic and nuclear localization. While transgenic lines GME-GFP, VTC4-GFP and l-GalDH-GFP driven by their endogenous promoters accumulated the fusion proteins, the functional VTC2-GFP protein is detected at low level using immunoblot in a complemented *vtc2* null mutant. This low amount of VTC2 protein and the extensive analyses using multiple combinations of SW enzymes in *N. benthamiana* supported the role *of VTC2* as the main control point of the pathway on ascorbate biosynthesis. Interaction analysis of SW enzymes using yeast two hybrid did not detect the formation of heterodimers, although VTC1, GME and VTC4 formed homodimers. Further coimmunoprecipitation (CoIP) analysis indicted that consecutive SW enzymes, as well as the first and last enzymes (VTC1 and l-GalDH), associate thereby adding a new layer of complexity to ascorbate biosynthesis. Finally, metabolic control analysis incorporating known kinetic characteristics, showed that previously reported feedback repression at the VTC2 step confers a high flux control coefficient and rationalizes why manipulation of other enzymes has little effect on ascorbate concentration.

**One sentence summary:** Metabolic engineering, genetic analysis and functional mutant complementation identify GDP-l-galactose phosphorylase as the main control point in ascorbate biosynthesis in green tissues.

## INTRODUCTION

l-Ascorbate (ascorbate, vitamin C) is the most abundant soluble antioxidant in plants and functions as a cofactor in many enzymatic reactions (Fenech et al., 2019; Smirnoff, 2018). Some groups of animals, among which humans are included, lost the ability to biosynthesize ascorbate (Smirnoff, 2018). Therefore, we need to incorporate this essential nutrient through the diet, with fruits and vegetables being the major source. In plants, ascorbate is synthesized from the hexose phosphate pool *via* d-mannose/l-galactose through the Smirnoff-Wheeler (SW) pathway (Wheeler et al., 1998; Fig. 1). The pathway is supported by abundant biochemical and genetic studies (Bulley and Laing, 2016; Wheeler et al., 2015). d-Glucose 6-P is sequentially transformed into d-fructose 6-P, d-mannose 6-P and d-mannose 1-P by phosphoglucose isomerase (PGI), phosphomannose isomerase (PMI) and phosphomannomutase (PMM). GDP-d-mannose pyrophosphorylase (VTC1; GMP) then converts d-mannose 1-P into GDP-d-mannose, which is subsequentially epimerized by GDP-d-mannose epimerase (GME) into GDP-l-galactose. GDP-d-mannose and GDP-l-galactose are additionally used in synthesis of cell wall polysaccharides and protein glycosylation thereby interconnecting the SW pathway with cell wall biosynthesis (Conklin et al., 1999; Fenech et al., 2019; Reiter and Vanzin, 2001). The next step is catalyzed by GDP-l-galactose phosphorylase (GGP), producing l-galactose 1-P. This step is proposed to be the main control point of the pathway (Dowdle et al., 2007; Laing et al., 2015, 2007; Linster et al., 2007; Yoshimura et al., 2014). In *Arabidopsis thaliana* (Arabidopsis), GGP activity is encoded by two paralogues, *VTC2* and *VTC5*. Next, l-galactose 1-phosphate phosphatase (VTC4, GPP) and other unidentified phosphatases (Conklin et al., 2006; Torabinejad et al., 2009), transform l-galactose 1-phosphate into l-galactose. l-Galactose is then oxidized to l-galactono-1,4-lactone by a l-galactose dehydrogenase (l-GalDH). All these steps are believed to occur in the cytosol while l-galactono-1,4-lactone crosses the outer membrane of the mitochondria to be oxidized by l-galactono-1,4-lactone dehydrogenase (l-GalLDH) which is an integral component of mitochondrial electron transport Complex 1 on the inner membrane (Pineau et al., 2008; Schimmeyer et al., 2016).

**Figure 1.**
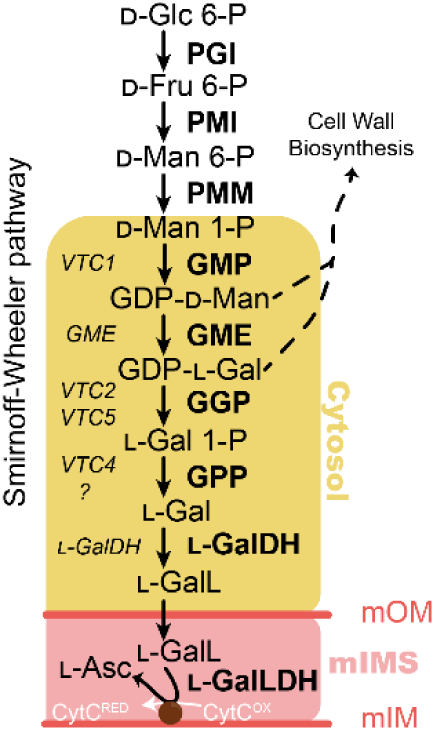
The ascorbate biosynthesis *via* GDP-mannose and l-galactose (the Smirnoff-Wheeler [SW] pathway) in *Arabidopsis thaliana*. Yellow area refers to the cytosolic enzymes used in this work. mOM: mitochondrion outer membrane, mIMS: mitochondrion intermembrane space, mIM: mitochondrion inner membrane. In the left-hand side of the arrows, the genes encoding each enzyme are displayed while in the right-hand side the encoded proteins/enzymatic activities are shown. Glc=glucose, Fru=fructose, Man=mannose, Gal=galactose, GalL=galactono-1,4-lactone, Asc= ascorbic acid. PGI=phosphoglucose isomerase, PMI=phosphomannose isomerase, PMM=phosphomannomutase, GMP=GDP-Mannose pyrophosphorylase, GME=GDP-mannose 3’,5’ epimerase, GGP= GDP-l-galactose phosphorylase, GPP= l-galactose 1-phosphate phosphatase, l-GalDH= l-galactose dehydrogenase, l-GalLDH= l-galactono-1,4-lactone dehydrogenase. Cyt^RED^=cytochrome c reduced, Cyt^OX^=cytochrome c oxidized.

Ascorbate concentration is dependent on the tissue (Franceschi and Tarlyn, 2002; Lorence et al., 2004; Müller-Moulé, 2008; Zhang et al., 2011) and the ascorbate pool adjusts to changes in environmental conditions, particularly light (Bartoli, 2006; Page et al., 2012; Plumb et al., 2018). Surprisingly, despite being the most abundant soluble antioxidant, a good understanding of the mechanisms that control ascorbate concentration remains limited. While mutant analyses of genes involved in biosynthetic pathway have corroborated the involvement of the various enzymes (Table 1), they provide little information on how the pathway is controlled. This is further complicated because the ascorbate pool is determined by the balance between synthesis, oxidation, recycling and breakdown. The oxidation product of ascorbate is the monodehydroascorbate radical. This is either directly reduced back to ascorbate or can give rise to dehydroascorbate (DHA) in its hydrated bicyclic hemiketal form, which is reduced back to ascorbate through the Halliwell-Foyer-Asada cycle (Asada, 1999). In addition, a proportion of ascorbate and DHA can be degraded to various products such as oxalate, threonate, and tartrate (Debolt et al., 2007; DeBolt et al., 2006; Dewhirst et al., 2017; Green and Fry, 2005; Pallanca and Smirnoff, 2000; Terai et al., 2020; Truffault et al., 2017). These recycling and breakdown reactions are evident under severe oxidative stress, which results in decreased cellular ascorbate (Terai et al., 2020; Waszczak et al., 2018).

**Table 1.** Phenotypes associated to Arabidopsis mutants affected in the genes involved in ascorbate biosynthesis available in the literature. Phenotype analysis shows that VTC2 is the first enzyme fully dedicated to ascorbate synthesis since growth arrest occurring in the *vtc2*/*vtc5* double mutant and downstream mutants is rescued by ascorbate supplementation. Although a reduced function (knock-down mutation) of genes upstream of GGP (*VTC2, VTC5)* also results in lower ascorbate concentration, the lethality occurring in knock-out mutants cannot be prevented by exogenous ascorbate supplementation.

| Enzyme | Mutant | Allele | Phenotype | Reference |
| --- | --- | --- | --- | --- |
| GMP | *cyt1-1* | Knock-out | Embryo lethal | Lukowitz et al., 2001 |
|  | *vtc1-1* | Knock-down | 30% of WT ascorbate | Conklin et al., 2000, 1996 |
| GME | *gme-1* | Knock-out | Male gametophyte lethal. Growth defect, rescued by boron but not by ascorbate supplementation. | Qi et al., 2017 |
|  | *gme-2* | Knock-down | 30% of WT ascorbate |  |
| GGP | *vtc2-4* | Single knock-out | 20% of WT ascorbate | Lim et al., 2016 |
|  | *vtc5-2* | Single knock-out | 80% of WT ascorbate | Dowdle et al., 2007; Lim et al., 2016 |
|  | *vtc2/vtc5* | Double knock-out | Growth arrest, rescued by ascorbate supplementation |  |
| GPP | *vtc4-4* | Knock-out | 65% of WT ascorbate | Torabinejad et al., 2009 |
| L-GalDH | *lgaldh* | Not reported | 30% of WT ascorbate in antisense suppression lines | Gatzek et al., 2002 |
| L-GalLDH | *gldh* | Knock-out | Growth arrest, rescued by ascorbate supplementation | Pineau et al., 2008 |

Multiple lines of evidence point to VTC2 as the critical controlling step of ascorbate biosynthesis. The transcription of *VTC2* gene (Dowdle et al., 2007; Müller-Moulé, 2008; Urzica et al., 2012) and the activity of the encoded enzyme (Dowdle et al., 2007)are highly responsive to environmental factors. Furthermore, *VTC2* translation is subject to feedback repression *via* a conserved upstream open reading frame (uORF) in the 5’-UTR (Laing et al., 2015). As a consequence, removing the uORF results in increased ascorbate in Arabidopsis (Laing et al., 2015), tomato (Li et al., 2018) and lettuce (Zhang et al., 2018). In addition, *VTC2* is the only gene of the pathway whose overexpression consistently increases the ascorbate content (Bulley et al., 2012; Yoshimura et al., 2014) and QTL analysis shows that *GGP* paralogs located within regions of the genome associated to fruits containing high ascorbate content (Mellidou et al., 2012). There have been reports that overexpression of other genes of the pathway such as *VTC1* or *GME* also increase ascorbate concentration in Arabidopsis (Zhou et al., 2012 [1.3-fold]; Li et al., 2016 [1.5-fold]), rice (Zhang et al., 2015 [1.4-fold]), tobacco (Wang et al., 2011 [2-4-fold]), tomato (Cronje et al., 2012 [1.7-fold]; Zhang et al., 2011 [1.4-fold]) and kiwifruit (Bulley et al., 2009 [1.2-fold]). However, overexpression of these genes does not always have an effect on ascorbate concentration (Sawake et al., 2015; Yoshimura et al., 2014). In addition, various studies have reported ascorbate feedback inhibition of the biosynthetic enzymes PMI (Maruta et al., 2008), GME (Wolucka and Van Montagu, 2003) and l-GalDH (Mieda et al., 2004). This feedback control is further supported by the decreased incorporation of labelled sugars into ascorbate *in vivo* after feeding ascorbate (Pallanca and Smirnoff, 2000; Wolucka and Van Montagu, 2003).

In order to increase our understanding of the organization of the ascorbate biosynthesis pathway, we have performed a systematic over-expression of single and multiple combinations of all the enzymes of the pathway from VTC1 downstream in *Nicotiana benthamiana*. This allowed the systematic investigation of their effects on the ascorbate pool and supported the formation of a multiprotein complex. In addition, complementation of the Arabidopsis mutants with the respective GFP-tagged enzymes confirmed the functionality of the constructs and provided information on their subcellular localizations. Finally, we constructed a kinetic model based on the known properties of the pathway whose predictions support that activity of VTC2 is the only significant controlling step and which also explains other observed features of the pathway. In addition to this information, this work provides a set of tools that will allow a better understanding of ascorbate biosynthesis at the molecular and cellular level.

## RESULTS

### Generation and analyses of GFP-tagged ascorbate biosynthesis Arabidopsis complementation lines reveal major differences in protein expression along the pathway

In order to study the function of SW proteins we aimed to generate Arabidopsis lines transformed with genomic regions (including their promoters) of ascorbate biosynthesis genes that resulted in proteins fused to GFP at their C-terminus. To investigate the functionality of these constructs, they were introduced into Arabidopsis mutants and the complementation of their phenotypes was analyzed. Arabidopsis mutants in SW genes typically show lower ascorbate concentration and/or lethality in early stages of development (Table 1). Selfing of Arabidopsis heterozygous mutants for *GME* (SALK_150208, *gme-3*, hereafter *gme*; Supp. Fig. 1A) and *l**-GalDH* (SALK_056664, Supp. Fig. 1B) did not produce homozygous mutants (Supp. Fig. 2). Since exogenous supplementation of ascorbate prevented growth arrest of several SW mutants (Dowdle et al., 2007; Pineau et al., 2008) we germinated seeds from heterozygous *gme-3* and *gldh* plants on solid media without and supplemented with 0.5 mM ascorbate (Fig. 2A). As a control we included seeds from a heterozygous *gldh* mutant plant, whose growth arrest is prevented by exogenous ascorbate (Pineau et al., 2008). The analysis of heterozygous *gme* progeny identified 40 wild-type (WT) seedlings, 46 heterozygous seedlings and no homozygous for the T-DNA, confirming the lethality of a loss-of-function allele of *GME* (Qi et al., 2017; Voxeur et al., 2011) (Fig. 2A; Supp. Fig. 2A). Similarly, no homozygous *gme* seedlings were identified when the growth medium was supplemented with 0.5 mM ascorbate in contrast to *gldh* (Pineau et al., 2008; Supp. Fig. 2B), confirming that GME, similarly to VTC1 (Lukowitz et al., 2001), has additional roles to that of ascorbate biosynthesis (Supp. Fig. 2A; Gilbert et al., 2009; Qi et al., 2017; Voxeur et al., 2011). On the other hand, diagnostic PCR of heterozygous *lgaldh* progeny identified 23 WT seedlings and 69 heterozygous T-DNA seedlings while no homozygous mutants were found in the medium not supplemented with ascorbate. Remarkably, a number of seedlings showed growth arrest and yellowing similarly to *gldh* progeny (Fig. 2A, magnified square). In contrast, we identified 25 WT, 41 heterozygous T-DNA and 25 homozygous T-DNA seedlings by diagnostic PCR in ascorbate containing medium (Fig. 2A, Supp. Fig. 2B). Therefore, and in contrast to *gme* mutants, *Igaldh* mutants could be rescued by supplementation with exogenous ascorbate (Fig. 2A, Supp. Fig. 2B). This supports that the expression product of *l**-GalDH* is responsible for the l-galactose dehydrogenase activity present in Arabidopsis. Additionally, the finding that, like *gldh, Igaldh* is an ascorbate auxotroph also indicates that the activity of this enzyme is essential for ascorbate biosynthesis.

**Figure 2.**
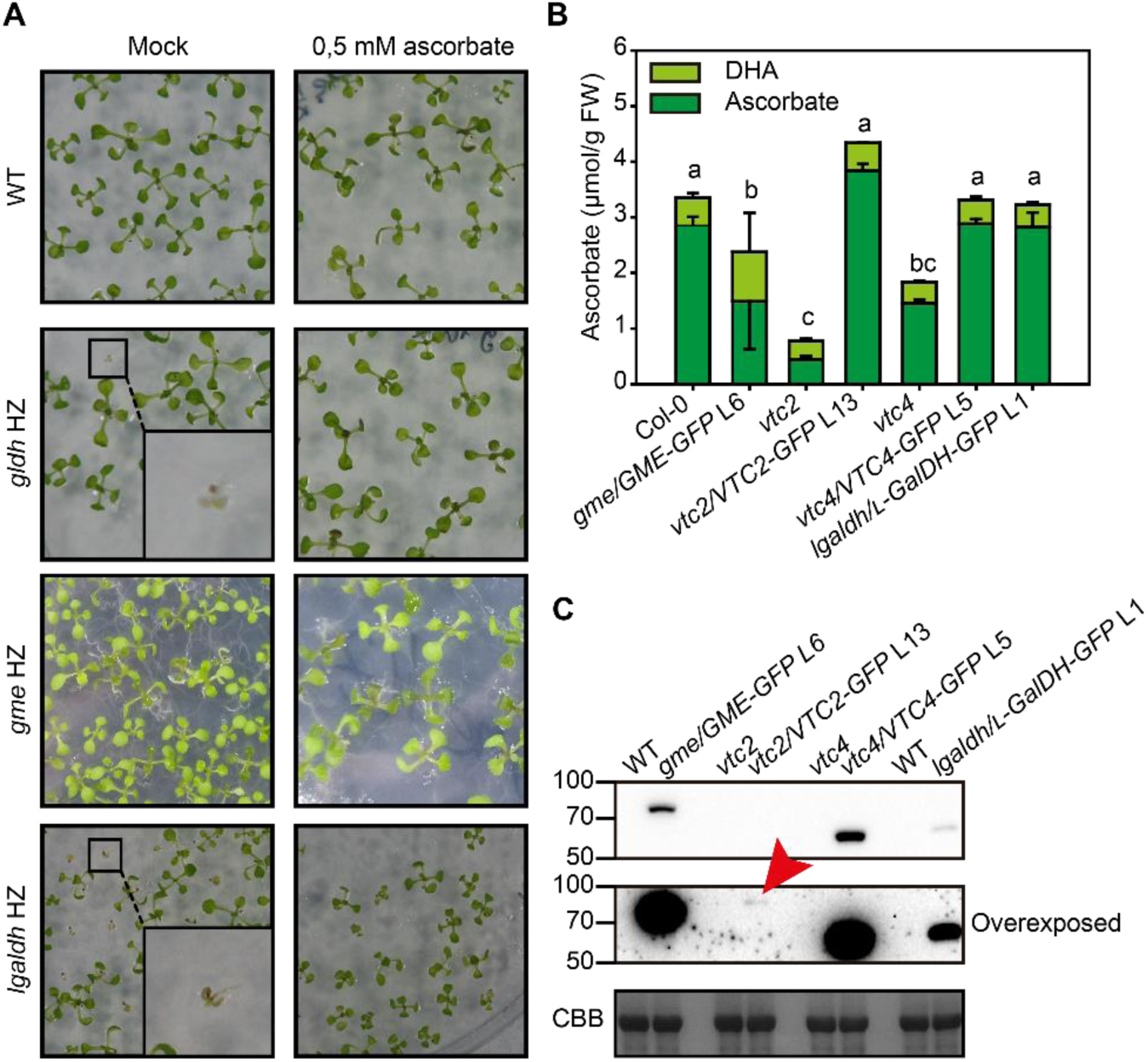
Characterization of C-terminal GFP fusions of ascorbate biosynthesis enzymes. A) Ascorbate complementation assay rescues growth arrest from *gldh* and *lgaldh* mutants but not *gme* lethality. In small dashed circles, homozygous mutants show arrested growth in media lacking ascorbate, but not in media supplemented with ascorbate. B) Complementation of ascorbate concentration by expression of GFP fusion proteins in ascorbate deficient Arabidopsis mutants. Different letters denote statistically significant differences for ascorbate concentration (One-Way ANOVA, α=0,05). Ascorbate was extracted from three independent samples composed of three 6-weeks old fully expanded rosettes grown in short day regime at 150 µmol photons m^-2^ s^-1^, one and a half hours after lights turned on. C) Immunoblot (α-GFP) of complemented lines show that little amount of protein VTC2 compared other components restores ascorbate content. Red arrowhead indicates VTC2-GFP. A representative sample for each genotype is shown out of the three tested by immunoblot (not shown).

In order to investigate the functionality of a GME protein fused with a GFP at the C-terminus, we transformed WT plants with a *GMEp:GME-GFP* construct. Of the several transformant that showed high expression of GME-GFP and a single insertion (Supp. Fig. 3A) line 6 (L6) was selected to introduce the transgene into the *gme* mutant background using reciprocal crosses. Using heterozygous *gme* plants as male parent, we did not obtain F1 *gme/GMEp:GME-GFP* (hereafter named *gme*/*GME-GFP* L6) descendants, supporting the role of *GME* in pollen development and pollen tube elongation (Qi et al., 2017). However, when line L6 was used as a male parent we obtained viable *gme/GME-GFP* F1 seedlings that allowed the identification in the F2 of homozygous *gme/GME-GFP* L6 seedlings (Supp. Fig. 3B). Surprisingly, while the GME-GFP fusion protein complement the sterility defects, it contained ∼55% of the ascorbate content of WT plants (Fig. 2B) which, considering the lower ascorbate content of knock-down *gme* Arabidopsis mutants (Qi et al., 2017), indicates that it complements only partially the ascorbate content. Similarly, a *l**-GalDHp:**l**-GalDH-GFP* construct was transformed into WT plants and selected line 1 (L1) that showed high expression and a single insertion (Supp. Fig. 3C). Then, reciprocal crosses between heterozygous *lgaldh* plants and *l**-GalDHp:**l**-GalDH-GFP* L1 were performed. In this occasion, we did obtain F1 plants when *lgaldh* plants were used as male or female parent. Homozygous *lgaldh/GalDH-GFP* L1 seedlings were identified in F2 progeny using diagnostic PCR in medium lacking ascorbate (Supp. Fig. 3D), indicating that the l-GalDH-GFP protein driven by the *l**-GalDH* promoter restores the enzyme activity lost in *lgaldh* mutant. In addition, ascorbate content of *lgaldh/**l**-GalDH-GFP* L1 plants contained WT levels of ascorbate (Fig. 2B).

In contrast to *gme* and *lgaldh*, loss of function *vtc2* and *vtc4* homozygous mutants are viable and show WT growth at standard conditions despite presenting a 20% and 55% of WT ascorbate content, respectively (Fig. 2B; Conklin et al., 2000; Dowdle et al., 2007; Torabinejad et al., 2009). In the case of *vtc2* mutant, GDP-l-galactose phosphorylase activity is also provided by the paralogous *VTC5* gene. Consistently, *vtc2vtc5* double mutants, like *lgaldh* and *gldh* mutants (Fig. 2A; Supp. Fig. 2B, C), show growth arrest that can be complemented with exogenous ascorbate (Dowdle et al., 2007; Lim et al., 2016). However, considering the phenotypes described (Table 1) the viability of *vtc4* must rely on other enzymes with l-galactose 1-phosphate phosphatase activity (Nourbakhsh et al., 2015; Torabinejad et al., 2009). In order to test the functionality of GFP-tagged VTC2 and VTC4, we relied on restoring WT ascorbate content by introducing *VTC2p:VTC2-GFP* and *VTC4p:VTC4-GFP* constructs into their respective mutants. T-DNA knock-out mutant *vtc2-4* (SAIL_769_H05; hereafter named *vtc2*; Supp. Fig. 1C; Lim et al., 2016) was directly transformed and lines with single insertion were selected (Supp. Fig. 3E). Among these lines, line 13 (L13) was selected because it showed the highest accumulation of VTC2-GFP (Supp. Fig. 3F). Also, *VTC4p:VTC4-GFP* was introduced into *vtc4-4* mutant (Supp. Fig 2D) following the same strategy described for *GME* and *l**-GalDH* (Supp. Fig. 3G). A single insertion *VTC4p: VTC4-GFP* line 5 was selected to cross with *vtc4* homozygous mutant and the homozygous *vtc4/VTC4-GFP* line was isolated by diagnostic PCR (Supp. Fig. 3H). Ascorbate concentration of *vtc2/VTC2-GFP* and vtc4/VTC4-GFP lines was restored to WT levels validating the functionally of the GFP-tagged enzyme (Fig. 2B).

We analyzed the protein levels of the different lines generated in a single blot in order to compare the amount of proteins accumulated in each line. Immunoblot analysis indicated that GME-GFP and VTC4-GFP were easily detectable while l-GalDH-GFP showed reduced levels of proteins (Fig. 3C). Interestingly, despite the complementation of ascorbate levels in L13, VTC2-GFP protein was not detected (Fig. 2C, upper panel) and was detected only after overexposure of the blot using a high sensitivity ECL substrate for the detection of femtogram amounts of protein. This low amount of VTC2 protein cannot be explained by a low expression of *VTC2* at the transcriptomic level because, based on public available transcriptomic databases, *VTC2* and *GME* show the highest expression of all SW genes in vegetative tissue (Supp. Fig. 4).

**Figure 3.**
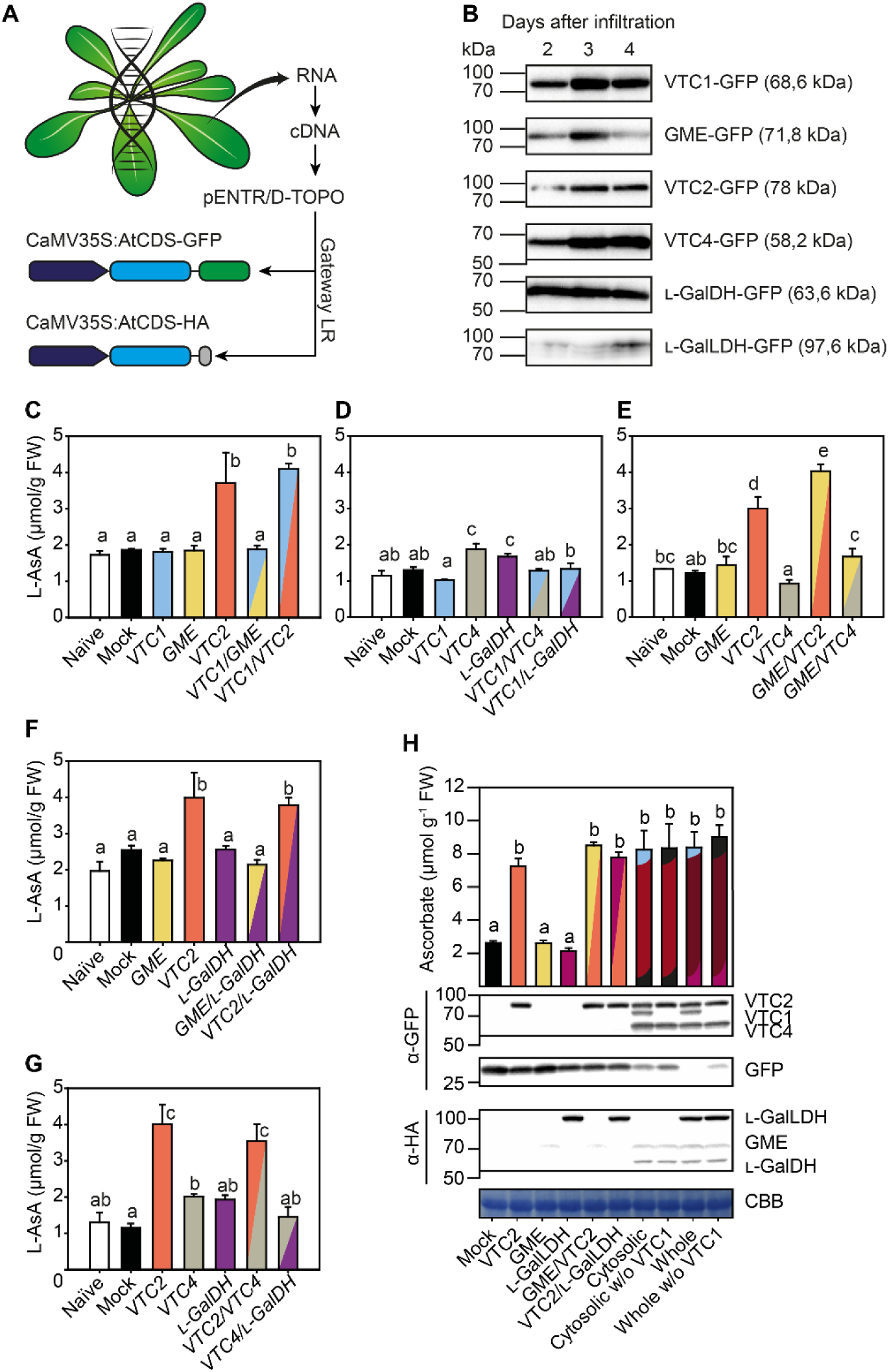
The effect of transient expression of C-terminal GFP and HA fusions of ascorbate biosynthesis enzymes on ascorbate concentration in *Nicotiana benthamiana* leaves. A) Strategy followed to clone and to overexpress in *N. benthamiana* leaves ascorbate biosynthesis genes from Arabidopsis translationally fused to GFP and HA at the protein C-terminus. B) Immunoblots (α-GFP) of fusion protein accumulation 2, 3 and 4 days after agroinfiltration. Expression was driven by the 35S promoter. Protein amount was measured and loaded to obtain similar intensities in the blot at 2 days after infiltration. C-H) Leaf ascorbate concentration 3 days after agroinfiltration with different combinations of C-terminal GFP ad HA fusions of ascorbate biosynthesis enzymes. Constructs used in each infiltration are shown in Supplementary Table 1. Two leaves of at least three *N. benthamiana* plants were infiltrated and samples were collected at 3 days after infiltration.

### Combinatorial expression of ascorbate biosynthesis genes in *N. benthamiana* leaves identifies VTC2 as the main control point of ascorbate biosynthesis

We have shown that that SW proteins are functional when a tag is added to their C-terminus and that a small amount of VTC2-GFP protein is required to increase the content of *vtc2* ascorbate to WT levels. In order to investigate the effect of the activity of ascorbate biosynthesis genes on ascorbate concentration we cloned the coding sequences (CDS) of Arabidopsis *VTC1, GME, VTC2, VTC4*, *l**-GalDH* and *l**-GalLDH* with C-terminal GFP and HA, driven by the CaMV35S promoter (Fig. 3A) and transiently expressed them in *N. benthamiana* leaves. Immunoblot analysis of the SW GFP tagged proteins showed that the molecular masses of the different fusion proteins were consistent with their predicted sizes (Fig. 3B). Moreover, all the fusion proteins showed little degradation of the proteins at all time points tested (Supp. Fig. 5). Three days after agroinfiltration, VTC1-GFP, GME-GFP, VTC2-GFP and VTC4-GFP proteins showed the highest expression while l-GalDH-GFP and l-GalLDH-GFP were also well expressed (Fig. 3B).

Then, we performed a systematic analysis of the effect of transient overexpression of individual or multiple combinations of these *SW* Arabidopsis genes on the ascorbate concentration in *N. benthamiana* leaves. Based on our previous data (Fig. 3B), we analyzed the ascorbate content 3 days after agroinfiltration. All genes were expressed either individually or in pairs (Fig. 3C-G; Supp. Table 1), and due to the large number of possible combinations, three genes were expressed individually and two genes out of the three were co-expressed in order to analyze the ascorbate content in the same experiment (Fig. 3C-G). This experimental design validated the reproducibility of our experimental system since individual genes have been analyzed three times with their proper controls (except for *VTC1*, with only two). As shown in Fig. 3C, the transient expression of individual *VTC1-GFP* and *GME-GFP* genes did not increase ascorbate content relative to the naïve (not agroinfiltrated) or control (infiltrated with GFP) *N. benthamiana* leaves. However, expression of *VTC2-GFP* increased ascorbate concentration between 2 and 3-fold. The expression of individual *VTC1, GME* and *l**-GalDH* genes or any combination of these three genes without *VTC2-GFP* did not significantly increase ascorbate concentration in any of the multiple assays (Fig. 3D-G). It also demonstrates that higher levels of VTC1 (Fig. 3C), VTC4 (Fig. 3G) and l-GalDH (Fig. 3F) do not contribute to increase the ascorbate content, either individually or combined. The combination of *VTC2* with *GME* caused a significant increase in ascorbate relative to *VTC2* (Fig. 3E), although the effect was not consistent in all experiments (Fig. 3E, H). An increased ascorbate content by combining *GME* and *VTC2* has been previously reported using transient overexpression of kiwifruit *GME* and *VTC2* homologs in tobacco leaves (Bulley et al., 2009).

Finally, we tested the effect of the simultaneous transient overexpression of all SW cytosolic Arabidopsis genes on the ascorbate concentration in *N. benthamiana* leaves. The expression of all cytosolic components was confirmed by immunoblot analysis (Fig. 3H) and increased the ascorbate content to similar levels as produced by expressing *VTC2* alone (Fig. 3H). Expressing all the cytosolic enzymes together with the final mitochondrial enzyme *l**-GalLDH* also did not increase ascorbate concentration (Fig. 3H). Overall, these results point to *VTC2* as the main control point in the biosynthesis of ascorbate.

### Subcellular localization of ascorbate biosynthesis enzymes using GFP-tagged proteins

The biosynthesis of ascorbate up to the l-galactose dehydrogenase step is proposed to take place in the cytosol with the exception of the last step that occurs at the intermembrane space of the mitochondria (Dunkley et al., 2006; Heazlewood et al., 2004; Østergaard et al., 1997; Pineau et al., 2008; Schimmeyer et al., 2016). *In silico* analysis using the software *Compartments* (https://compartments.jensenlab.org/Search) predicted a cytosolic subcellular localization for GME, VTC4 and l-GalDH, a dual cytosolic-nuclear localization of VTC1 and VTC2 and a mitochondrial localization of l-GalLDH (Supp. Fig. 6). Confocal microscopy imaging of C-terminal GFP-tagged proteins transiently expressed in *N. benthamiana* revealed a cytosolic subcellular localization for VTC1, GME, VTC2, VTC4 and l-GalDH while l-GalLDH-GFP appeared in structures that resembled mitochondria (Fig. 4). Interestingly, VTC1-GFP, VTC2-GFP, VTC4-GFP and l-GalDH-GFP also showed a nuclear localization in addition to their cytoplasmic localization, whereas we observed a GME-GFP signal surrounding the nuclei (Fig. 4). Evidence for a dual cytosolic-nuclear localization has been provided for VTC1 and VTC2 using stable transgenic Arabidopsis lines (Müller-Moulé, 2008; Wang et al., 2013), suggesting a potential nuclear function of these proteins. Remarkably, similarly to VTC1, nuclear localization signals are not predicted for VTC4 or l-GalDH and, therefore, the localization reported here was unexpected (Supp. Table 2).

**Figure 4.**
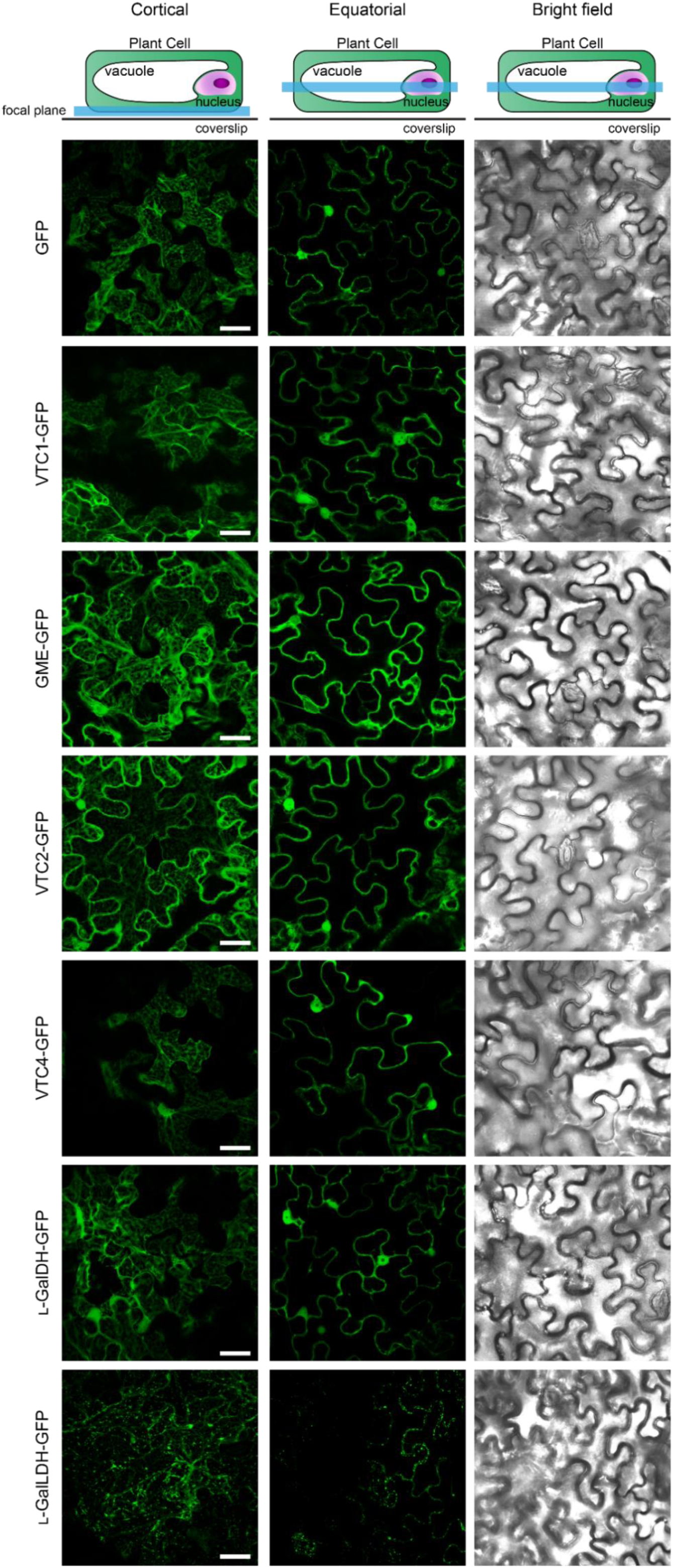
Subcellular localization of free GFP and C-terminal GFP fusions of ascorbate biosynthesis enzymes transiently expressed in *Nicotiana benthamiana* leaves under the control of the 35S promoter 3 days after agroinfiltration. GFP was visualized by laser scanning confocal microscopy. Scale bar = 30 µm.

Because overexpression can result in ectopic localization of proteins, we used the Arabidopsis transgenic lines that carry functional biosynthetic ascorbate proteins C-terminally fused to GFP and driven by their own promoters to analyze their *in vivo* subcellular localization as well as their tissue distribution. Cotyledons, cotyledon-hypocotyl junction, hypocotyl and root tip of four-day-old Arabidopsis seedlings were analyzed. GME-GFP, VTC4-GFP and l-GalDH-GFP lines showed a strong fluorescence signal in the cytoplasm (Fig. 5A). In addition, VTC4-GFP and l-GalDH-GFP were present within nuclei-like structures. In roots, these two proteins were present within nucleus as a dot that resembles nucleolus, but they also displayed a ring-like pattern in nucleus whereas GME-GFP showed an even distribution across cytosol (Fig. 5A). We did not detect fluorescence signal for VTC2-GFP above the background level (Fig. 5B) consistent with the weak signal previously observed in immunoblot analysis (Fig. 2H). Except for VTC2, these data are consistent with the over-expression data of *SW-GFP* constructs in *N. benthamiana*.

**Figure 5.**
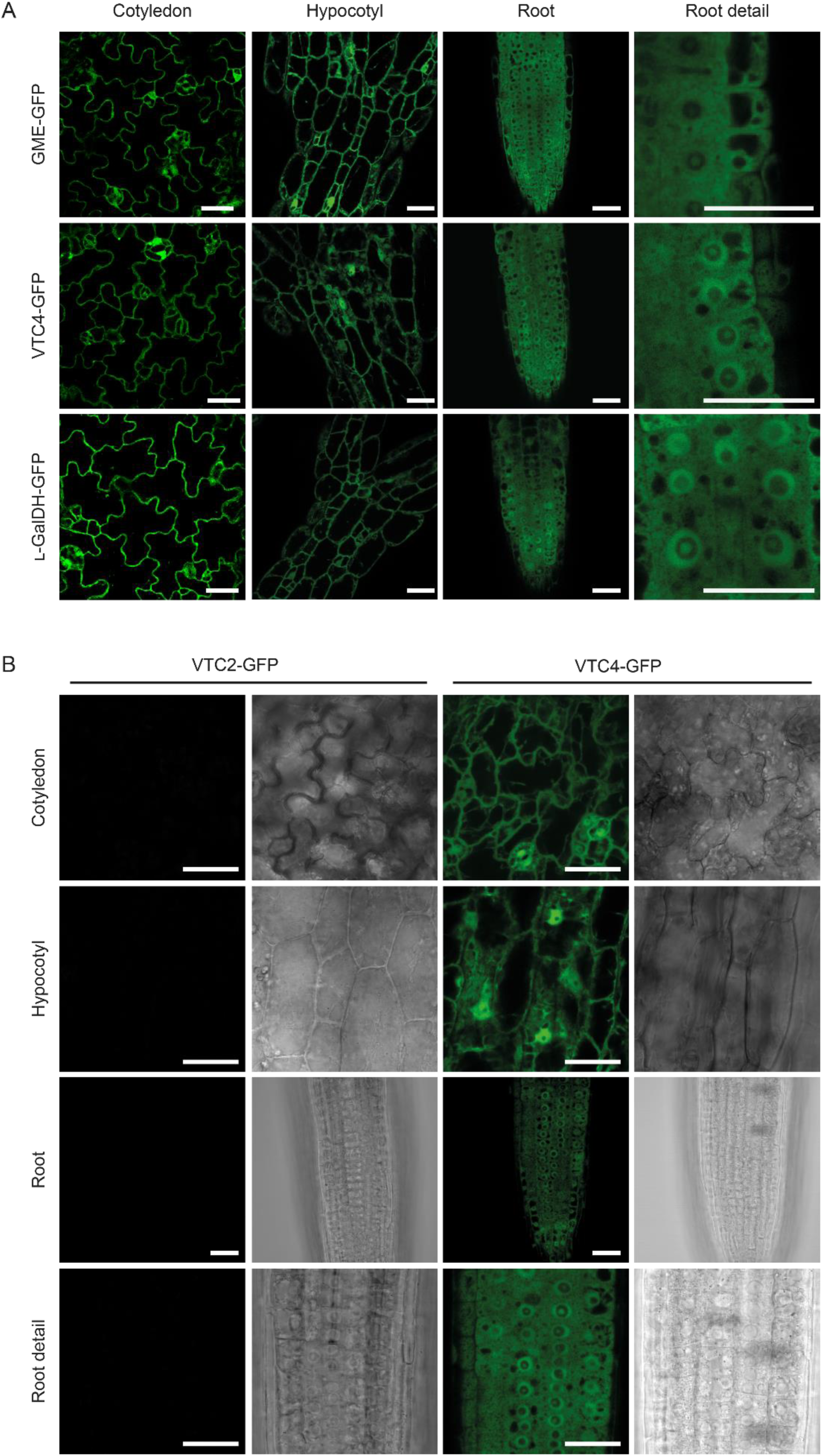
Subcellular localization of ascorbate biosynthesis enzymes fused to GFP at the C-terminus and expressed under the control of their native promoters in Arabidopsis. A) The expression of GFP-tagged enzymes is ubiquitous in four days-old seedlings and their subcellular localization is compatible with cytosol and nucleus except for GME, which only locates in cytosol. B) GFP signal of a functional VTC2-GFP (line 13) protein was not detectable. GFP was visualized by laser scanning confocal microscopy. Scale bar = 30 µm.

### Protein-protein interaction assays support a physical association of proteins involved in ascorbate biosynthesis

Enzymes involved in metabolic pathways can associate in order to increase the efficiency of consecutive enzymatic transformations (Smirnoff, 2019; Sweetlove and Fernie, 2018). Since VTC1, GME, VTC2, VTC4 and l-GalDH proteins showed cytoplasmic localization (Fig. 4, Fig. 5) we investigated their interaction *in viv*o using yeast two-hybrid (Y2H) (Fig. 6A; Supp. Fig. 7). For this, all ascorbate biosynthesis CDS (including the mitochondria-localized l-GalLDH as a negative control) were cloned into both binding domain (BD, pGBKT7) and activation domain (AD, pGADT7) yeast vector and tested all 36 possible interactions resulting from the combination of all the enzymes of the pathway in both directions (fused to AD or to BD; Supp. Fig. 7). As positive controls, we included SNF1/SNF2 (Fields and Song, 1989) and p53/SV40-Large AgT (Iwabuchi et al., 1993; Li and Fields, 1993) interactions, whereas Laminin C/SV40-Large AgT was used as a negative control (Bartel et al., 1993; Ye and Worman, 1995). This analysis did not provide evidence of direct interactions between the enzymes of the pathway, although VTC1, GME and VTC4 showed self-interaction, showing that these proteins were expressed in yeast (Fig. 6A). A dimerization interface is predicted for GME and VTC4, while a trimerization domain is predicted for VTC1 (Fig. 6B), consistent with our yeast two-hybrid interaction results.

**Figure 6.**
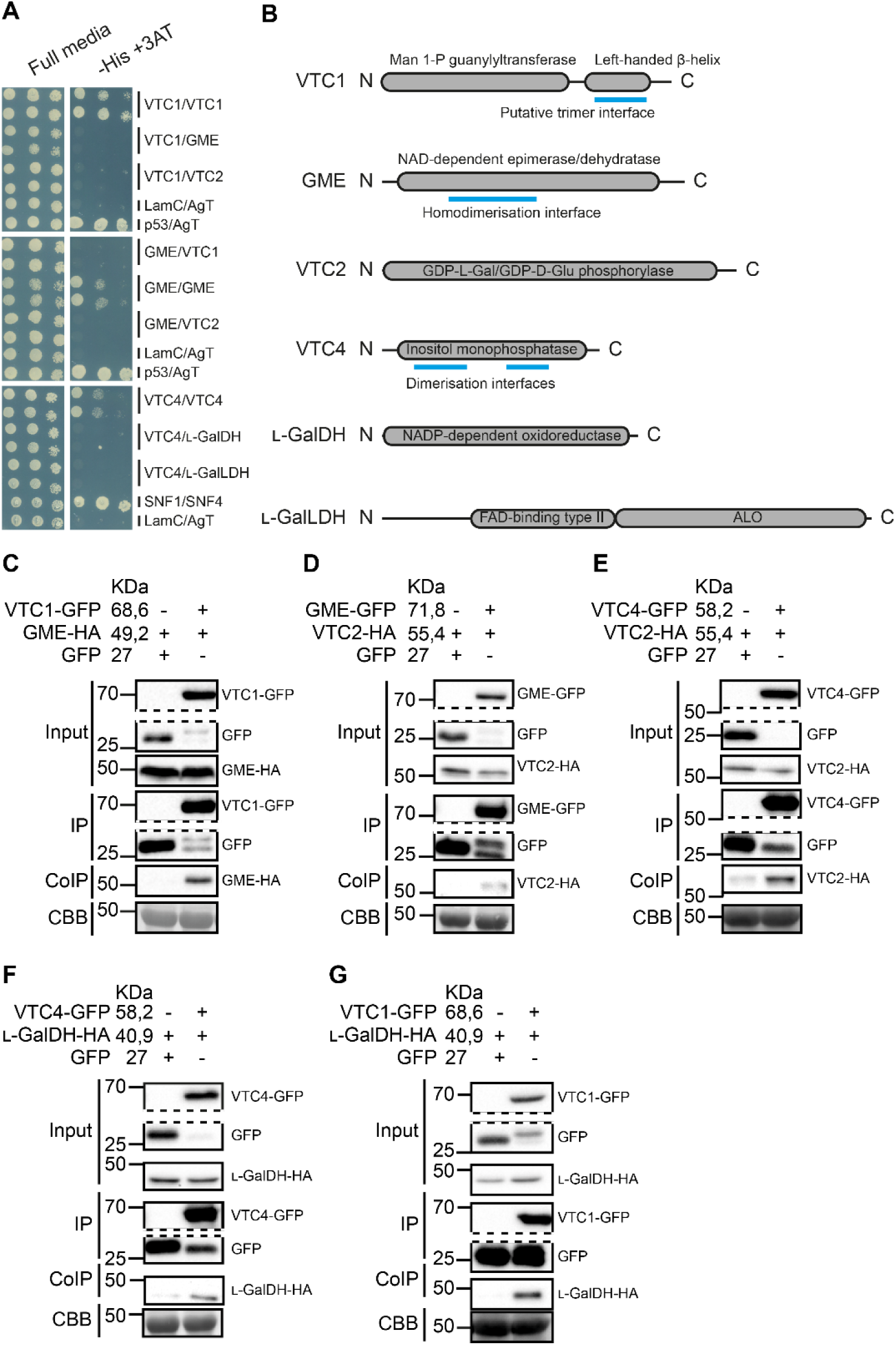
Interaction of ascorbate biosynthesis enzymes assessed by yeast two-hybrid analysis and coimmunoprecipitation (CoIP). (A) Yeast two-hybrid assay detected no direct interaction between the enzymes, but VTC1, GME and VTC4, which contain predicted dimerization/trimerization domains (B), form dimers/trimers. B) Protein scheme for each of the enzymes assayed in yeast two-hybrid showing length (in amino acids), annotated domains and predicted dimer/trimerization interfaces. (C-G) CoIP assay reported association in vivo in Nicotiana between consecutive steps as well as between the first (VTC1) and the last (l-GalDH) steps occurring in cytosol. GFP-tagged enzyme from Nicotiana crude protein extracts containing two overexpressed consecutive enzymes, GFP and HA-tagged, was pulled-down using α-GFP agarose beads and coimmunoprecipitated protein was detected using α-HA antibody.

Despite the apparent lack of direct interactions of ascorbate biosynthesis enzymes in yeast, we investigated whether these proteins could associate *in vivo*. For this, we co-expressed biosynthesis enzymes of the pathway tagged with C-terminal GFP and HA in *N. benthamiana* and performed coimmunoprecipitation (CoIP) assays. We used GFP-Trap beads to inmunoprecipitate GFP tagged proteins and α-HA antibodies to detect co-immunoprecipitated HA-fused proteins using immunoblot. We first tested biosynthesis proteins that are consecutive in the pathway (Fig. 6C-F), i.e., VTC1/GME, GME/VTC2, VTC2/VTC4, VTC4/l-GalDH. We co-expressed VTC1-GFP and GME-HA to test the interaction, and GFP together with GME-HA as a negative control (Fig. 6C) and checked by immunoblot if protein expression was suitable for immunoprecipitation prior to the assay (Supp. Fig. 8). After immunoprecipitation of VTC1-GFP and free GFP, we detected a specific association between GME-HA with VTC1-GFP but not with free GFP (Fig. 6C). Following the same strategy, we found associations between all consecutive enzymes of the pathway (Fig. 6D-F). In addition, we also performed CoIP between the first and last cytosolic enzymes of the pathway, i.e., VTC1 and l-GalDH and found a specific association between VTC1-GFP and l-GalDH-HA (Fig. 6G).

### A kinetic model of ascorbate biosynthesis indicates GDP-l-galactose phosphorylase as the main control point

To further understand the results of over-expression, a kinetic model of ascorbate biosynthesis was constructed using Complex Pathway Simulator (COPASI; Hoops et al., 2006). Reactions were described by reversible Michaelis-Menten kinetics, except for L-Gal 1-P phosphatase/VTC4 and L-GalLDH which were described by irreversible Michaelis-Menten kinetics (Fig. 7A). Published Km values were used except for GGP, which was set to 0.1 mM in the absence of published data (Gatzek et al., 2002; Hoeberichts et al., 2008; W. A. Laing et al., 2004; Linster et al., 2007; Maruta et al., 2008; Østergaard et al., 1997; Wolucka and Van Montagu, 2003). The relative activity of enzymes has been reported in only one case (Dowdle et al., 2007) but these are not guaranteed to reflect the maximum rate of each enzyme. Low expression of GGP compared to other pathway enzymes is suggested from very low expression of the GFP fusion which nevertheless rescues ascorbate concentration in the mutant background (Fig. 3D, H). However, conservatively, the relative activity of each enzyme (V) was set to the same value (10 mM s-1) but was varied during simulations. Evidence from radiolabeling suggests that ascorbate synthesis is subject to feedback inhibition by ascorbate (Conklin et al., 1997; Laing et al., 2015; Pallanca and Smirnoff, 2000; Wolucka and Van Montagu, 2003). The proposed uORF-mediated feedback repression of GGP translation (Laing et al., 2015) was modelled by including non-competitive inhibition by ascorbate. Competitive inhibition of PMI and l-GalDH by ascorbate (Maruta et al., 2008; Mieda et al., 2004) were included, although in the case of l-GalDH, the inhibition could be an assay artefact (William A Laing et al., 2004). Once synthesized, ascorbate and its oxidation product dehydroascorbate (DHA) can be broken down to a wide range of products (Dewhirst et al., 2020; Dewhirst and Fry, 2018; Green and Fry, 2005). The oxidation of ascorbate to monodehydroascorbate (MDHA) and DHA was included in simplified form including only ascorbate and DHA. The breakdown rate of DHA was described by irreversible 1st order kinetics (Conklin et al., 1997; Pallanca and Smirnoff, 2000). It is assumed in the model that all breakdown products are recycled to d-Glc 6-P. This is unlikely to be correct but did not affect model predictions.

**Figure 7.**
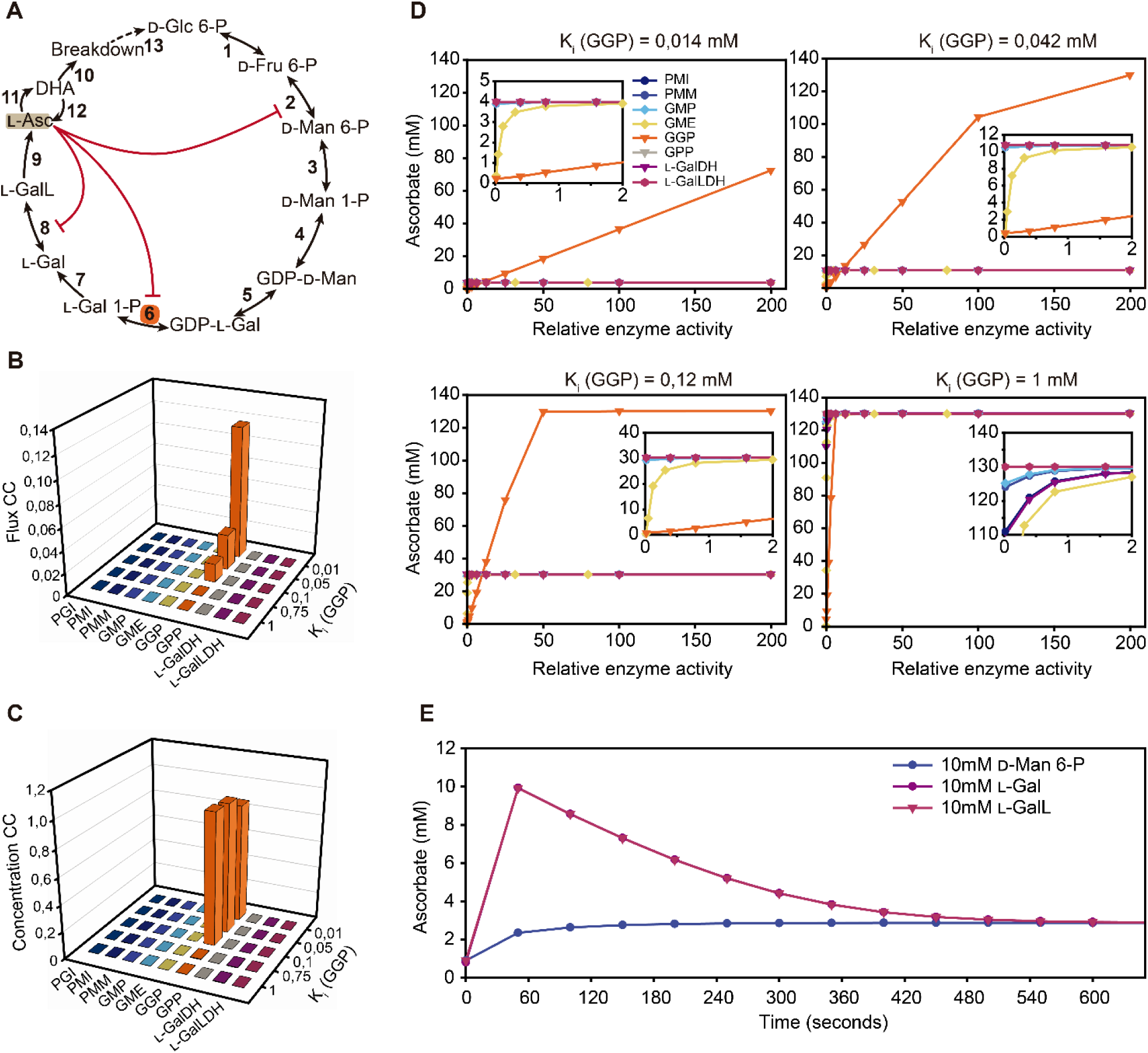
A kinetic model of ascorbate biosynthesis and turnover. A) Scheme of the pathway modelled in COPASI showing intermediates catalytic steps (arrows) and feedback (red lines). The initial model parameters are shown in Supplementary Table 3. B) Metabolic control analysis at steady state was used to calculate flux control coefficients for each enzyme with various strengths of non-competitive inhibition exerted by ascorbate (indicated by K_i_ values) on GGP (step 6). C) Metabolic control analysis at steady state was used to calculate concentration control coefficients for ascorbate with various strengths of non-competitive inhibition exerted by ascorbate (indicated by K_i_ values) on GGP (step 6). D) The effect of varying enzyme activity on ascorbate concentration with various strengths of non-competitive inhibition exerted by ascorbate (indicated by K_i_ values) on GGP (step 6). E) Time course of the change in ascorbate concentration in response to the addition of d-mannose 6-phosphate (d-Man 6-P), l-galactose (l-Gal) and l-galactono-1,4-lactone (l-GalL). 1: PGI, 2: PMI, 3: PMM, 4: GMP, 5: GME, 6: GGP, 7: GPP, 8: l-GalDH, 9: l-GalLDH, 10: Turnover, 11: Oxidation, 12: Reduction).

The starting parameters used in the model are shown in Supplementary Table 3. The model carried out metabolic control analysis (MCA) at steady state. The effect of changing the strength of feedback repression of GGP by ascorbate on the flux control coefficients for each step are shown in Figure 7B (and data in supplemental Table 4). At the highest Ki value (relatively weak feedback), all steps up to GGP contributed equally to flux, indicated by similar flux control coefficients. However, biosynthesis steps beyond GGP had very low flux control coefficients and changes in activity of these enzymes would be predicted to have a very small effect on pathway flux. The turnover step also had a significant effect on flux. Decreasing Ki by ten-fold or more resulted in all flux control being shared between GGP and turnover, the pre-GGP steps having very low flux control coefficients. Therefore, for the biosynthesis steps, the model predicts that only GGP controls the flux through the pathway if feedback repression is sufficiently strong. Correspondingly, concentration control coefficients (i.e. the change in the concentration of each pathway intermediate caused by a change in enzyme activity) for each step were calculated. The results for ascorbate are shown in Figure 7C and for all intermediates in Supplementary Table 5. At the highest Ki value, all steps up to GGP had similar concentration control coefficients indicating change in activity at each step would exert a similar effect on ascorbate concentration. Increasing feedback strength decreased the concentration control coefficients of all steps except GGP predicting that only this step controls ascorbate concentration. The outcome could be affected by the starting conditions (relative enzyme activities and kinetic constants) but testing the model with varying Km values or with and without competitive feedback inhibition of PMI and l-GalDH had little effect on ascorbate concentration (Supplementary Table 6). To simulate metabolic engineering experiments, the enzyme activity of each step was increased over a large range and the predicted change in ascorbate concentration at various feedback repression strengths was determined (Fig. 7D and supplementary Table 7). As expected from the MCA, only varying GGP activity had a significant effect on ascorbate over a very wide range of activities and this effect was strongly dependent on the extent of feedback repression. In comparison, it was necessary to decrease the activity of the other enzymes to very low values before ascorbate concentration was affected and this behavior was particularly marked for the post-GGP steps. The kinetic model therefore reflects the effect of transient expression of the pathway enzymes in N. benthamiana: only increasing GGP activity increases ascorbate concentration. Simulation of mannose 6-P addition caused a small transient increase in ascorbate accumulation, while l-galactose addition had a large effect (Fig. 7E). This differential response reflects the operation of feedback repression at the GGP step.

## DISCUSSION

Here, we report the effect of systematic overexpression, individually and in combination, of all genes involved in the SW pathway have on the endogenous ascorbate content using *N. benthamiana* leaves. The GFP-tagged enzymes were functional since translational fusion of GFP at the C-termini of all enzymes increased ascorbate concentration in their respective Arabidopsis mutants. Further, immunoblots of the expressed enzymes showed that they were all well expressed in *N. benthamiana* (Supp. Fig. 8). Under these experimental conditions, only VTC2/GGP over-expression was able to increase ascorbate concentration, which confirms previous studies (Ali et al., 2019; Bulley et al., 2012, 2009; Laing et al., 2015; Li et al., 2018; Yoshimura et al., 2014; Zhang et al., 2018). Interestingly, a vtc2 mutant containing a VTC2-GFP construct driven by its own promoter and including the 5’-UTR showed an extremely low GFP fluorescence compared to other SW GFP-tagged proteins. This low expression of VTC2-GFP was also reflected in the immunoblot comparing the expression of all the components of the pathway at the protein level (Fig. 2E), likely reflecting the translational repression of VTC2 via the conserved uORF in the 5’-UTR (Laing et al., 2015). This is further confirmed by the finding that the overexpression in *N. benthamiana* of VTC2 without the uORF showed a similar amount of VTC2 proteins to other proteins of the pathway (Supp. Fig. 8). The expected low level of endogenous VTC2 protein in *N. benthamiana*, which also contains the uORF (Laing et al., 2015), may explain the marked increase of ascorbate after the overexpression of Arabidopsis VTC2 lacking the uORF.

Based on the data presented here and other reports, we constructed a kinetic model based on the known properties of the pathway enzymes that includes the repression of GGP by ascorbate. However, it does not explicitly distinguish between transcriptional, translational, post-translational and metabolite feedback on enzyme activity as the mechanism of repression. This model was used for metabolic control analysis, an approach which determines the distribution of the control of flux and concentration of pathway intermediates in response to small changes in activity of each enzyme in the pathway by calculating flux control and concentration control coefficients for each step (Fell, 1992). This analysis showed that the only step in the biosynthesis pathway that controls pathway flux and ascorbate pool size is GGP as long as feedback repression is included. The other critical step is ascorbate breakdown while the flux control coefficients of all the other steps were negligible. Decreasing the strength of feedback repression by ascorbate on GGP eventually decreased its flux and concentration control coefficients, resulting in control being more evenly distributed across the steps up to GGP. The model predicts that the steps post GGP exert little control and that in the presence of feedback repression, GGP is the only critical step. Consequently, in virtual metabolic engineering experiments, in which each enzyme was changed over a large range, only GGP affected ascorbate concentration.

The model prediction is consistent with the transient expression results shown in this report and with the relatively small effect of over-expressing enzymes other than GGP. In a previous study of transient expression in *Nicotiana tabacum*, GME alone did not affect ascorbate, in accordance with the model, but enhanced the effect of expressing GGP (Bulley et al., 2009). However, in our hands this increase in ascorbate was not always reproducible suggesting that other factors need to be considered such as different carbohydrate status caused by growth conditions. Stable over-expression of GMP and GME has been reported to increase ascorbate in tomato and *N. tabacum* (Badejo et al., 2008, 2007; Li et al., 2019) and the model predicts that these enzymes could be effective if endogenous activity is low. Similarly, increasing the expression of l-GalDH in *N. tabacum* had no effect on ascorbate while antisense suppression in Arabidopsis only had a small effect in high light (Gatzek et al., 2002).

The model outcomes are robust over a range of starting conditions and are minimally affected by the Km values tested nor by the presence/absence of competitive inhibition of PMI and l-GalDH. The model also explains another feature of the pathway, which is that VTC2 and VTC5, the paralogues of GGP in Arabidopsis, do not compensate for each other when knocked out (Dowdle et al., 2007). The control features of the pathway appear to emerge from its architecture in which a strongly repressible GGP step is followed by the irreversible l-Gal 1-P phosphatase and l-galactonolactone dehydrogenase steps.

In a previous study comparing the activity of SW enzymes where ascorbate pool size differs under low and high light intensity, only GGP showed increased activity in high light in Arabidopsis (Dowdle et al., 2007). Transcript levels suggest that only VTC2 and VTC5 (and sometimes GME) are responsive to environmental conditions (Li et al., 2013), and that there is not a coordinated induction of SW genes under high light and low temperature, conditions that increase ascorbate concentration (Laing et al., 2017). The control of ascorbate biosynthesis therefore differs from pathways in which there is coordinated induction of multiple pathway enzymes by transcription factors, for example anthocyanin synthesis (Zhang et al., 2014).

Importantly, the model shows that ascorbate pool size responds to GGP over a wide range and thereby explains how a combination of transcription and feedback repression controls the amount/activity of GGP and provide a mechanism to adjust ascorbate pool size to prevailing conditions. Light is the best studied factor and ascorbate adjusts to the prevailing light intensity over several days (Page et al., 2012). The mechanism likely involves a changed balance between increased transcription of GGP via a possible photosynthesis-derived signal and light response promoter elements (Gao et al., 2011; Yabuta et al., 2007) and ascorbate repression of translation via the uORF. In addition to the transient expression assays showing the role of the uORF in translational repression of GGP (Laing et al., 2015), CRISPR-Cas9 editing of the uORF in Arabidopsis, lettuce and tomato increases ascorbate, confirming its role in repression of ascorbate synthesis in diverse species (Li et al., 2018; Zhang et al., 2018). Faster control could also occur if ascorbate status more directly affects GGP activity via direct inhibition or post-translational modification. These latter possibilities have not been investigated but could operate since ascorbate addition relatively rapidly decreases incorporation of 14C-glucose or mannose into ascorbate (Pallanca and Smirnoff, 2000; Wolucka and Van Montagu, 2003). It is not known if ascorbate or a proxy of ascorbate concentration exerts repression and, also, the mechanism by which the uORF controls GGP translation is not known.

In addition to investigating the effect of enzyme overexpression on ascorbate accumulation, we used tagged enzymes to investigate subcellular localization and the formation of enzymatic complexes. With the exception of l-GalLDH, the cytosolic localization of the SW enzymes is consistent with the in-silico prediction (Supp. Fig. 6). In this study, we have shown that all SW proteins are present in the cytosol (Fig. 4; Fig. 5). Interestingly, in addition to VTC1 and VTC2 that were previously reported to localize in the nucleus (Müller-Moulé, 2008; Wang et al., 2013) VTC4 and l-GalDH also showed nuclear localization (Fig. 4; Fig. 5) despite the lack of nuclear localization signals (Supp. Table 2). Whether or not this nuclear localization is functionally important remains to be determined but several metabolic enzymes moonlighting in the nucleus have been previously reported (Boukouris et al., 2016). Using CoIP in *N. benthamiana* we have shown that enzymes catalyzing consecutive steps in the pathway associate (Fig. 6C-F) as well as proteins catalyzing distant enzymatic steps such as VTC1 and l-GalDH (Fig. 6G). However, no direct interactions were found using yeast two-hybrid analysis. The lack of direct interaction but the positive in vivo associations may indicate that the enzymes might attach to scaffold proteins so that the interaction would not be detectable in the yeast two-hybrid assay. Although this deserves further investigation it is tempting to speculate that some of the ascorbate biosynthesis enzymes might form a functional enzymatic complex (metabolon) that could potentially channel pathway intermediates to increase flux and perhaps aid partitioning of GDP-d-mannose and GDP-l-galactose between ascorbate and polysaccharide synthesis. Such channeling has been shown for a number of pathways (Amorim-Silva et al., 2019; Smirnoff, 2019; Sweetlove and Fernie, 2018; Zhang et al., 2017).

**In conclusion**, metabolic engineering experiments using the last six enzymes of the SW pathway show that only GGP has significant control over the pathway, and we have developed a kinetic model that provides a rationale for this result. It is clear that the balance of VTC2/VTC5 amount is controlled by a combination of transcription and translation repression, largely, but not entirely mediated by the uORF mechanism. In addition, we have produced a set of complemented mutants from GME to l-GalDH tagged with GFP that provides a resource for further investigation of the SW pathway.

## Materials and Methods

### Plant material

*Arabidopsis thaliana* (L.) Heynh wild-type (WT) and transgenic lines generated were in Col-0 ecotype. Arabidopsis mutants *vtc2-4* (SAIL_769_H05; Lim et al., 2016), *vtc4-4* (SALK_077222; Torabinejad et al., 2009) and *gldh* (SALK_060087; Pineau et al., 2008) used in this study have been previously described. However, to the best of our knowledge, this is the first time that SALK_150208 (here named *gme-3*) and SALK_056664 (here named *lgaldh-1*), obtained from the The European Arabidopsis Stock Centre (NASC: http://arabidopsis.info/), were described (M&M Supp. Table 1). Diagnostic PCR was performed to identify the presence of T-DNA insertion in individuals plants using the allele-specific primers listed in M&M Supp. Table 2A. The generation of stable transgenic lines either in WT or in the respective mutant backgrounds is described in the “Generation of transgenic lines” section and lines are listed in (M&M Supp. Table 1).

### Standard growth conditions

Arabidopsis seeds were surface sterilized using chlorine gas by pouring 3 mL of 37% HCl into 100 mL of commercial bleach and airtight sealed for 4 hours. Then, seeds were air-cleared for at least 4 hours in a laminar flow cabinet and stored at 4°C for 3-days stratification. Seeds were sowed on half-strength Murashige-Skoog (MS) agar-solidified (0.6% [w/v] agar) medium supplemented with 1,5% [w/v] sucrose under sterile conditions and grown with a long-day photoperiod (16-h light/8-h darkness cycle, 22±1 °C, 150±50 µmol photons m^-2^ s^-1^) for 7-10 days. Seedlings were then collected for analysis, transferred to liquid media or to soil (4:1 (v/v) soil:vermiculite).

### Plasmid Constructs

Two different types of DNA fragments were used as templates (genomic and coding DNA sequence) to generate the constructs used in this work. First, genomic fragments were amplified starting from at least 2 kb upstream the +1 ATG unless otherwise specified (Supp. Fig. 1) until the stop codon (not included) using genomic Col-0 DNA as template. Second, coding DNA sequences (CDS) were amplified, starting at +1 ATG until the stop codon (not included), using as template Col-0 Arabidopsis cDNA that was generated using total RNA. In the case of *VTC2*, due to a failure in amplifying the entire VTC2p:VTC2 product, about 1.4 kb (according to Gao et al., 2011) of the *VTC2* promoter including the 5’UTR was amplified (Supp. Fig. 1; M&M Supp. Table 3). The primers used to generate these above-mentioned DNA fragments are detailed in M&M Supp. Table 2B and 2C. A proofreading DNA polymerase was used for all these PCR amplifications (TAKARA PrimeSTAR Max DNA Polymerase, CAT. #R045A).

*VTC2* promoter was digested with HindIII and directly cloned into the *pGWB504* vector. All other DNA fragments (excluding *VTC2* promoter) were cloned into *pENTR/D-TOPO* using the pENTR Directional TOPO cloning kit (Invitrogen). Next, on the one hand, DNA genomic fragments (which includes promoter and lacks the stop codon) were recombined from *pENTR/D-TOPO* into *pGWB4* by LR reaction (Invitrogen) to generate *SWp:SW-GFP* constructs for *GME, VTC4* and *l**-GalDH*. On the other hand, all CDS fragments (without promoter, 5’-UTR and stop codon) were sub-cloned into *pGWB5* and *pGWB14* to generate *CaMV35S:SW_CDS_-GFP* and *CaMV35S:SW_CDS_-HA*, respectively. In the case of *VTC2*, the CDS was also recombined to *VTC2p-pGWB504* by LR reaction (Invitrogen) thus obtaining the *VTC2p:VTC2_CDS_-GFP* construct Likewise, CDS (*pENTR/D-TOPO*) were sub-cloned into both *pGADT7-GW* (activation domain, AD) and *pGBKT7-GW* (binding domain, BD) to generate *SW*_*CDS*_-*AD* and *SW*_*CDS*_-*BD*, respectively, for yeast two-hybrid assay (see Yeast two-hybrid assay section). The expression vectors generated using the destination vectors *pGWB5* and *pGWB14* contain a kanamycin and a hygromycin resistance gene for bacteria and plant selection, respectively. The yeast two-hybrid vectors *pGADT7* and *pGBKT7* contain an ampicillin and kanamycin resistance gene, respectively for selection in bacteria. A summary of generated constructs is showed in M&M Supp. Table 3. All constructs were analyzed by colony PCR (using the primers indicated in M&M Supp. Table 2D), enzymatic DNA restriction and sequencing (using the primers indicated in M&M Supp. Table 2E).

The plasmids used from the pGWB vector series were provided by Tsuyoshi Nakagawa (Department of Molecular and Functional Genomics, Shimane University; Nakagawa et al., 2007). The pGADT7(GW) and pGBKT7(GW) destination vectors were provided by Salomé Prat (Nacional de Biotecnología-Consejo Superior de Investigaciones Científicas).

### Generation of Arabidopsis transgenic lines

The constructs generated using *pGWB4* plasmid for native expression of the SW genes (pGWB504 for *VTC2*) were transformed into *Agrobacterium tumefaciens* GV3101::pMP90 by electroporation and confirmed by colony PCR using the primers indicated in M&M Supp. Table 2D. Arabidopsis WT (Col-0) plants were then transformed using these *A. tumefaciens* strains by floral dipping (Clough and Bent, 1998). T2 plants were analyzed for hygromycin resistance ratio resistant:sensitive (R:S) and lines with single insertion (T2 Hyg^R^:Hyg^S^=3, Χ^2^ (α=0.05), n>300) were selected. Selected T2 plants were grown in long day photoperiod and a sample of rosette leaf was taken to perform immunoblot analysis (Fig. 2; Supp. Fig. 3). Seeds from different T2 lines with a high expression were selected and homozygous T3 and T4 plants were used in this work.

For generation of the complementation lines, selected T2 plants (with single insertion) were crossed with their respective mutants in both directions as male and female. In order to obtain homozygous plants for both the T-DNA insertion and the SWp:SW-GFP construct (Hyg^R^), we selected hygromycin resistant plants and performed diagnostic PCR to identify T-DNA insertion using the primers in M&M Supp. Table 2A. Homozygous F3 plants for T-DNA and SWp:SW-GFP insertion were used in this work.

### Ascorbate complementation assay

WT, *gme* (SALK_150208), *lgaldh* (SALK_056664) and *gldh* (SALK_060087) seeds were surface sterilized and stratified for 3 days at 4 °C. Seeds were then sowed in agar-solidified (0.5% [w/v] agar) modified Scholl medium with or without 0,5 mM ascorbate as described elsewhere (Lim et al., 2016) and grown with a long-day photoperiod. After 10 days, plants were individually collected and PCR-diagnosed to identify homozygous mutants.

### Ascorbate measurements

#### High Performance Liquid Chromatography (HPLC) for combinatorial expression in Nicotiana benthamiana

Ascorbate was extracted from approximately 250 mg fresh tissue using 2 mL of a 2% (w/v) metaphosphoric acid, 2 mM EDTA ice-cold buffer (modified from Davey et al., 2006). Samples were centrifuged at 18000 g for 20 min at 4°C, and supernatants were filtered (0.45 µm) before injection into an HPLC (Agilent Technologies 1200 series; Rx-C18, 4.6×100 mm 3.5 µm column (Agilent Technologies 861967-902), diode array detector (Agilent Technologies G1315D). Samples were measured at 254 nm using filtered (0.45 µm) 0.1 M NaH2PO4, 0.2 mM EDTA, pH 3.1 (orthophosphoric acid) as mobile phase (Harapanhalli et al., 1993) at 4 °C (column at 20 °C; 5 µL sample injection, 0.7 mL/min flow, 200 bar). Standards were prepared using ascorbic acid dissolved in extraction buffer, with concentrations ranging from 0 mM to 2.5 mM.

#### Ascorbate oxidase (AO) activity for Arabidopsis transgenic lines

Ascorbate and dehydroascorbate were extracted from approximately 80 mg fresh tissue using ice cold 1 mL of a 3% (w/v) metaphosphoric acid plus 1 mM EDTA. Samples were centrifuged at 16000 g for 10 min at 4°C. Absorbance measurements were carried out at 265 nm using 20 µL sample/standard aliquots in a 96 well UV-transparent plate (Greiner UV-Star® 96-Well Microplate) mixed with 100 µL of either phosphate buffer (0.2 M KH2PO4, pH 7 to measure ascorbate) or Tris(2-carboxyethyl)phosphine (TCEP)-phosphate buffer (0.2 M KH2PO4, pH7, 2 mM TCEP, to measure ascorbate and dehydroascorbate). Then, 5 µL 40 U/mL AO was added to each well, mixed and incubated at room temperature for 20 minutes, when the absorbance was again measured. The difference in the absorbance was interpolated in the standard curve, using standards ranging from 0 mM to 1 mM in extraction buffer. TCEP and AO stocks were prepared in phosphate buffer.

### Immunoblot

Proteins were separated by size through a sodium dodecyl sulphate-polyacrylamide gel electrophoresis (SDS-PAGE) and transferred onto a polyvinylidene difluoride (PVDF) membrane (Immobilon-P, Millipore, 0.45 µm pore size; CAT. #IPVH00010) previously methanol-equilibrated. Next, the PVDF membrane was blocked with 5% (w/v) skimmed milk followed by an incubation with a primary antibody synthesized in mouse against GFP (Santa Cruz Biotechnology, CAT. #sc-9996; 1:600) or HA (Sigma-Aldrich, CAT. #H3663; 1:3000) epitopes. Then, the membrane was subsequentially incubated with a horseradish peroxidase (HRP)-conjugated α-mouse whole IgG antibody produced in rabbit (Sigma-Aldrich, CAT. #A9044; 1:80000). Protein-HRP chemiluminescence was detected using Chemidoc XRS1 System (Bio-Rad) after incubation with Clarity ECL Western Blotting Substrate or SuperSignal West Femto Maximum Sensitivity Substrate according to the manufacturer’s instructions.

### Transient expression in *Nicotiana benthamiana* leaves

*Agrobacterium tumefaciens* GV3101::pMP90 was transformed by electroporation with the constructs generated using pGWB5 and pGWB14 positive colonies were confirmed by colony PCR. Two leaves per plant (leaves 3 and 4 from apex) 4-week-old *N. benthamiana* were infiltrated in the abaxial side of using 1 mL syringe (without needle). Agrobacterium cultures were grown overnight in liquid Luria-Bertani medium containing rifampicin (50 μg/mL), gentamycin (25 μg/mL), and the construct-specific antibiotic kanamycin (50 μg/mL). Cells were then harvested by centrifugation (for 15 min at 3000 g in 50-mL falcon tubes) at room temperature, pellets were resuspended in agroinfiltration solution (10 mM MES, pH 5.6, 10 mM MgCl_2_, and 1 mM acetosyringone), and kept for 2-3 h in dark conditions at room temperature. In order to avoid overexpression-associated gene silencing, *Agrobacterium* strain containing a p19 expression vector was coinfiltrated. For single gene expression, the gene of interest and p19 were mixed in such a way that the optical densities (600 nm, OD_600_) were 0.8 and 0.2, respectively. For multiple co-infiltration experiments, the genes of interest were diluted in the same proportion so the final OD_600_ was 1 (0,8 for the combination of genes of interest and 0,2 for p19). Agrobacterium strains expressing either free GFP or TTL3-HA (an ascorbate biosynthesis non-related protein (Amorim-Silva et al., 2019)) were used to compensate OD_600_ in co-infiltration experiments.

To achieve optimal protein expression, immunoblot time course analysis was addressed 2, 3 and 4 days after infiltration (Fig. 3B). Protein loads were adjusted so comparable amounts of GFP-tagged protein were detected two days after infiltration for each construct using the Java-based image-processing program FIJI (Schindelin et al., 2012; Schneider et al., 2012).

### Confocal laser scanning microscopy

All confocal images of *N. benthamiana* (Fig. 4) were obtained using a Leica TCS SP5 II confocal microscope equipped with a 488-nm argon laser for GFP excitation and a PMT for its detection using the HCX PL APO CS 40×1,25 OIL objective with additional 2x digital zoom. Arabidopsis imaging was carried out using a Leica TCS SP8 equipped with HC PL APO CS2 40x/1,30 OIL objective and a solid state 488-nm laser used to excite GFP and a high-sensitivity SP hybrid detector (Fig. 5A) and a Zeiss LSM 880 (Fig. 5B) with a 488-nm argon laser and PMT detectors for GFP and transmitted light using the C-Apochromat 40x/1.2 W Korr M27 objective. All image processing was performed using FIJI (Schindelin et al., 2012; Schneider et al., 2012).

### Yeast two-hybrid assay

The Gal4-based yeast two-hybrid system (Clontech Laboratories) was used for testing the interaction. *Saccharomyces cerevisiae* Meyen ex E. C. Hansen strain PJ69-4A was transformed with the constructs described in the plasmid constructs section (SW genes clones into *pGADT7* and *pGBKT7*) as described elsewhere (Gietz and Schiestl, 1995). Constructs transformed are explained in the plasmid constructs section (*pGADT7* and *pGBKT7*). Transformants were grown on plasmid-selective medium (synthetic defined (SD)/ -Trp -Leu) and grown at 28 °C for 4 days. Two independent colonies were taken per transformation event and resuspended in 200 µL of sterile water. Ten-fold serial dilutions were made and 5 µL of each dilution were spotted onto three different interaction-selective medium: SD -Trp (minus tryptophan) -Leu (minus leucin) -His (minus histidine) +2 mM 3-AT (3-amino-1,2,4-triazole); SD -Trp -Leu -Ade; SD -Trp -Leu -Ade +3-AT. Full SD medium was also used to check strain survival. Plates were grown at 28 °C and pictures were taken after 3, 5 and 9 days (only 5 days is shown). Concentrations are available at Rodríguez-Negrete et al. (2014).

### Bioinformatic analyses

Protein domains predictions were performed using Protein Basic Local Alignment Search from the National Center for Biotechnology Information (Altschul et al., 1997, BLASTp; https://blast.ncbi.nlm.nih.gov/Blast.cgi?PAGE=Proteins) followed by functional annotation of protein domain using the Conserved Domain Database (Marchler-Bauer et al., 2017, 2015, 2011; Marchler-Bauer and Bryant, 2004; https://www.ncbi.nlm.nih.gov/Structure/cdd/cdd.shtml). Protein subcellular localization was predicted using COMPARTMENTS (Binder et al., 2014; https://compartments.jensenlab.org/Search). Predicted subcellular localization signals were performed using LOCALIZER (Sperschneider et al., 2017; http://localizer.csiro.au/). Metabolic control Analysis was carried out using Complex Pathway Simulator (COPASI, Hoops et al., 2006; http://copasi.org/). All the details needed for the simulations are collected in Supp. Tables 3-7). VTC2 mRNA expression datasets were extracted from Transcriptome Variation Analysis database (Klepikova et al., 2016, TRAVA; http://travadb.org/) and eFP-seq Browser (Sullivan et al., 2019; https://bar.utoronto.ca/eFP-Seq_Browser/).

### Coimmunoprecipitation assay

Four-week-old *N. benthamiana* plants were used for transient expression assays as described above. Leaves were ground to fine powder in liquid nitrogen. ∼0.5 g of ground leaves per sample was used, and total proteins were extracted with 1 mL of non-denaturing extraction buffer (50 mM Tris-HCl pH7.5, 150 mM NaCl, 1% (v/v) Nonidet P-40, 10 mM EDTA, 1mM Na2MoO4, 1 mM NaF, 10 mM DTT, 0.5 mM PMSF**, 1% (v/v) protease inhibitor (Sigma P9599)) and incubated for 30 min at 4 °C using an end-over-end rocker. Protein extracts were centrifuged at 20000 g for 20 min at 4 °C and then filtered by gravity using Poly-Prep chromatography columns (731-1550, Bio-Rad). Supernatants were filtered by gravity through Poly-Prep chromatography columns (731-1550, Bio-Rad), and 100 mL was reserved for immunoblot analysis as input. The remaining supernatants were used for immunoprecipitation of GFP-fused proteins agarose using GFP-Trap coupled to agarose beads (Chromotek) and following the manufacturer’s instructions. Total (input), immunoprecipitated (IP), and CoIP samples were resuspended with 2x concentrated Laemmli sample buffer and heated at 95°C for 5 min protein denaturation. Finally, samples were separated in a 10% SDS-PAGE gel and analyzed as described above.

## Statistical analysis

All analyses were performed using SigmaPlot v11 for Windows considering n>3 and α=0.05.

## Accession Numbers

Sequence data were sourced from the The Arabidopsis Information Resource (TAIR) (https://www.arabidopsis.org/) under the following accession numbers: AT2G39770 for *VTC1*, AT5G28840 for *GME*, AT4G26850 for *VTC2*, AT3G02870 for *VTC4*, AT4G33670 for *l**-GalDH*, and AT3G47930 for *l**-GalLDH*.

## Supplemental material

**Supplementary Figure 1.**
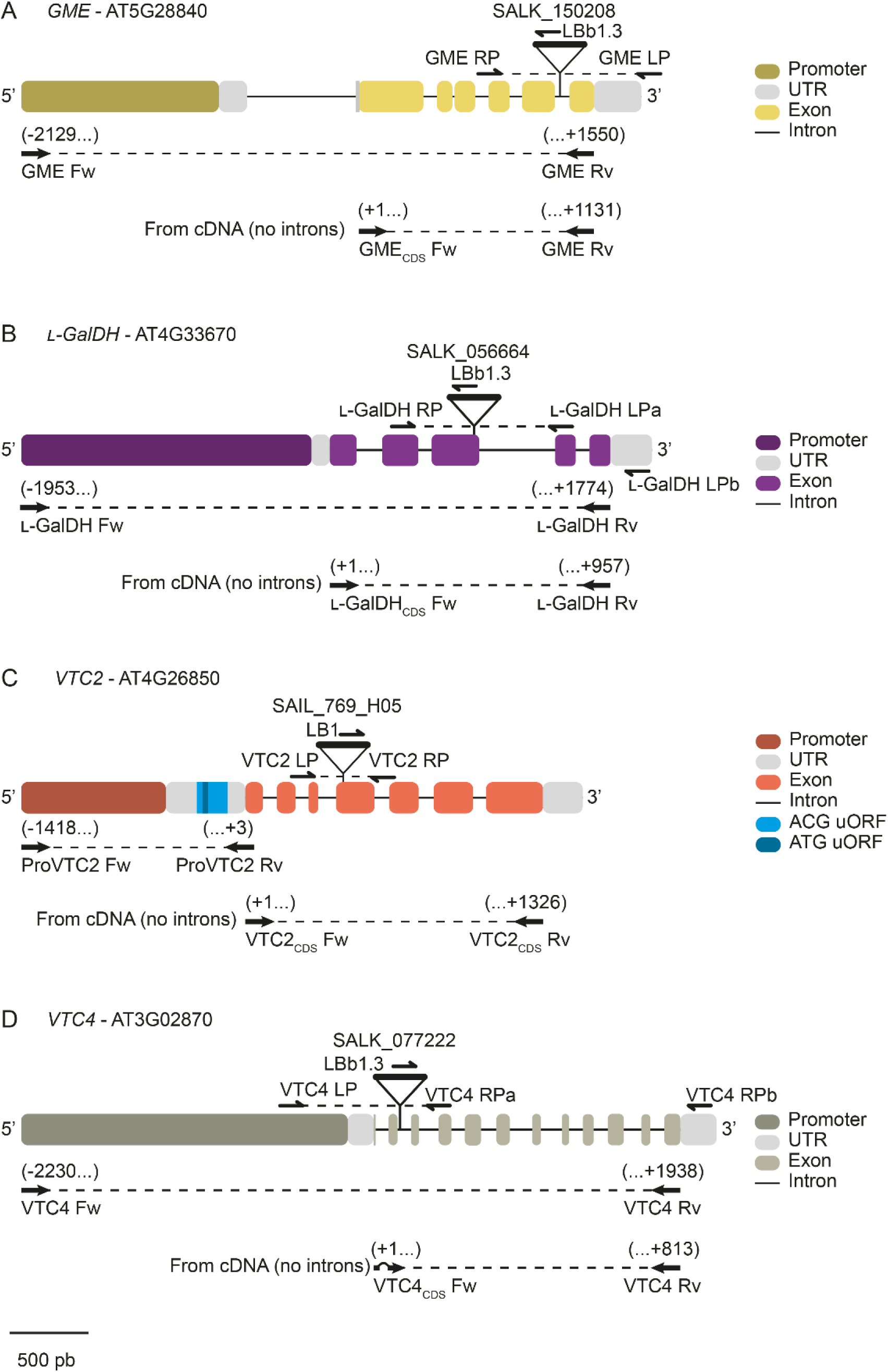
Gene structure, primers disposition and mutants’ T-DNA location.

**Supplementary Figure 2.**
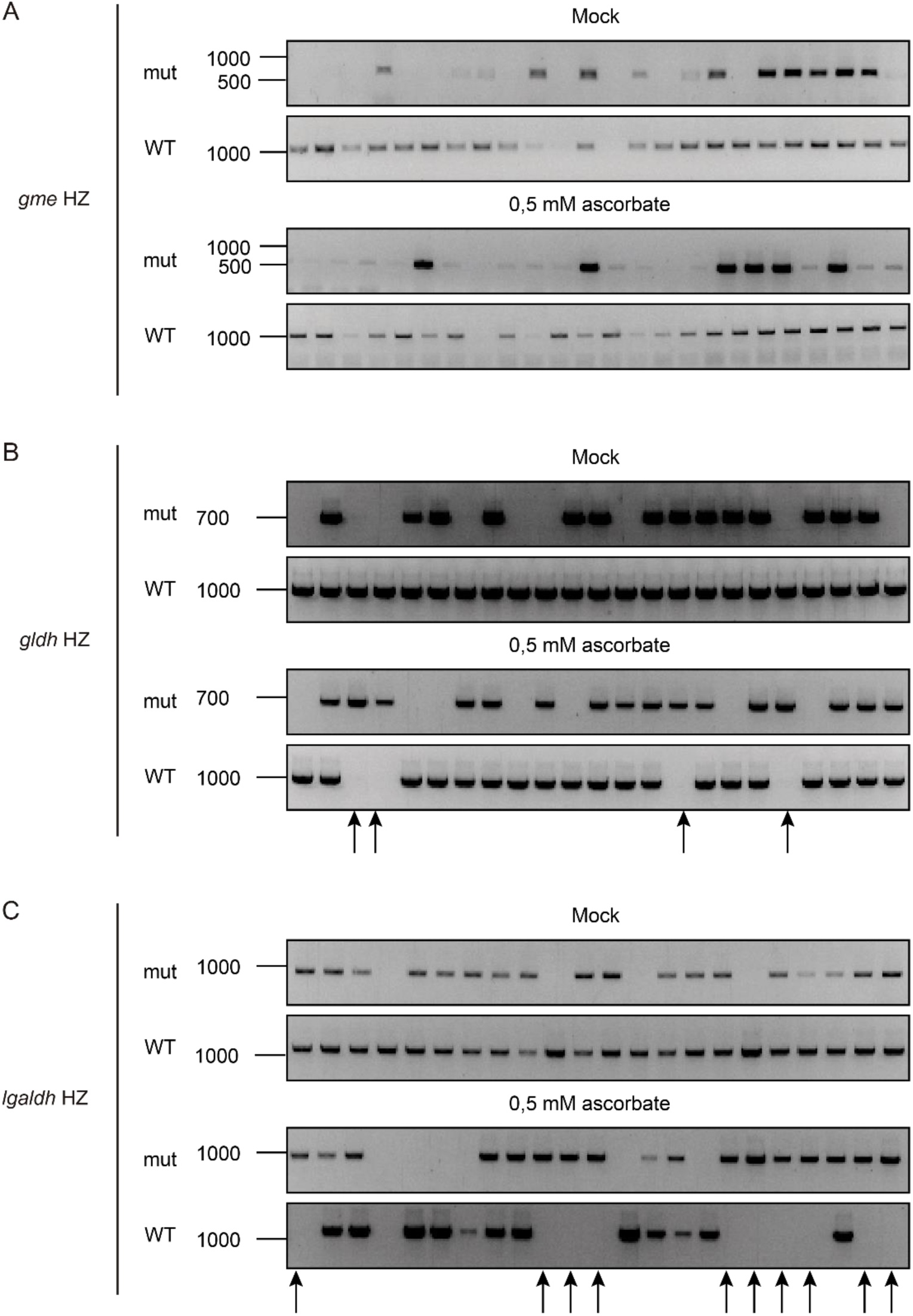
Diagnostic PCR of the *gme, lgaldh* and *gldh* seedlings analyzed in the complementation assay described in Figure 2A. Whereas ascorbate does not recover the lethality of *gme* mutation (A), it does recover *lgaldh* (C). As a control, we used *gldh* (B), previously published to be recovered by ascorbate (Pineau et al., 2008).

**Supplementary Figure 3.**
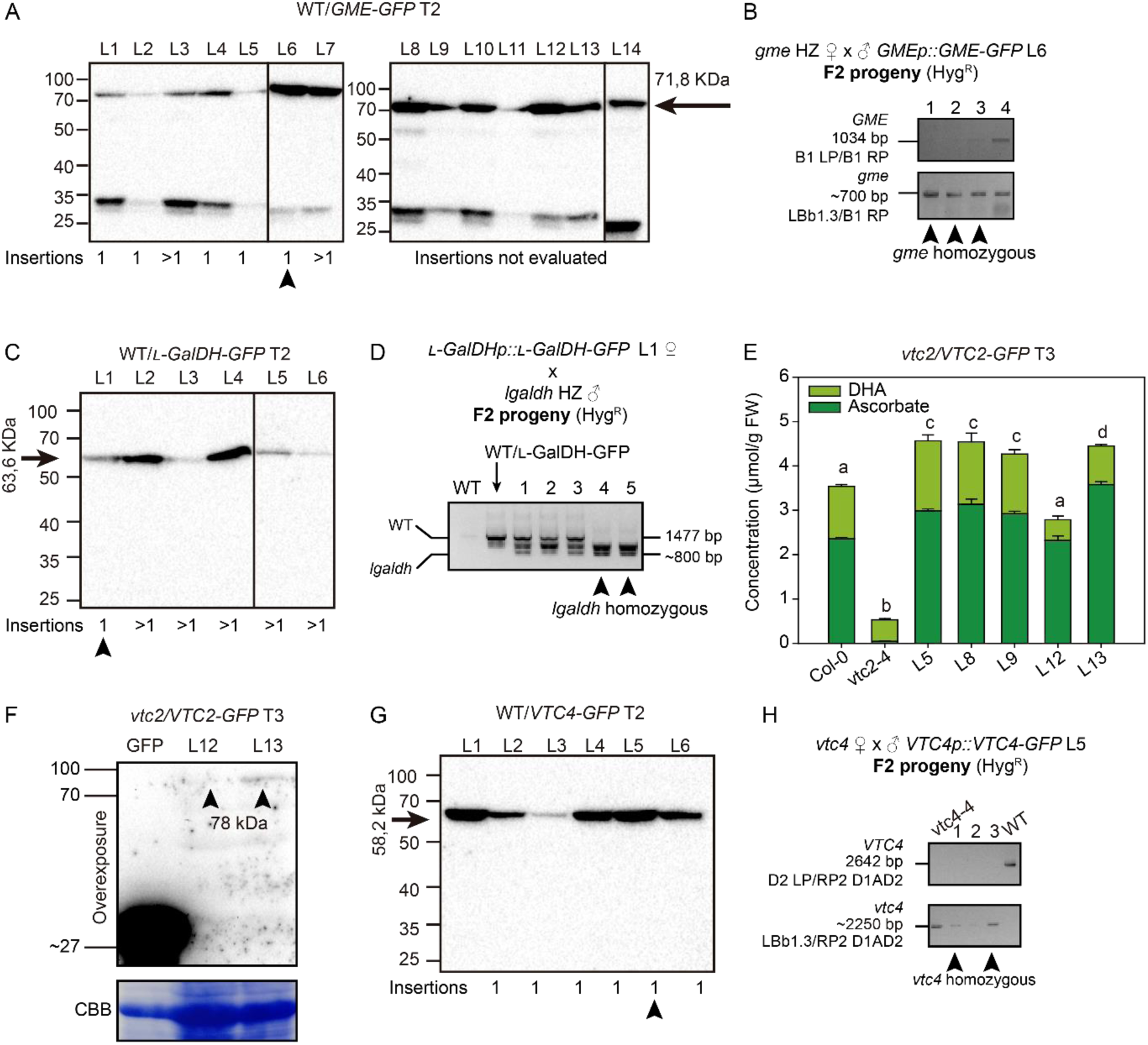
Generation of Arabidopsis transgenic lines containing ascorbate biosynthesis enzymes fused to GFP. A) Immunoblot (α-GFP) analysis of fourteen GME-GFP T2 lines identifies several lines expressing different amounts of GME-GFP protein (background WT). After the identification of lines harboring single insertions (T2 Hyg^R^:Hyg^S^=3, Χ^2^ (α=0.05)), we selected line L6 (arrowhead) to introduce the GMEp:GME-GFP construct into the *gme* mutant (SALK_150208) background using reciprocal crosses. B) Diagnostic PCR of F2 generation from the cross indicated identified plants harboring GME-GFP in homozygous mutant background (arrowheads). Ascorbate content of this line is shown in Figure 2. C) Immunoblot (α-GFP) analysis of VTC2-GFP of two lines among which ascorbate content ranges. Four days old seedlings were grown under long day conditions in solid half-strength MS agar plates. D) Immunoblot (α-GFP) analysis of l-GalDH-GFP T2 generation showing the six lines expressing l-GalDH-GFP protein (background WT). After the identification of lines harboring single insertions (T2 Hyg^R^:Hyg^S^=3, Χ^2^ (α=0.05)), we selected line L1 to introduce the l-GalDHp:l-GalDH-GFP construct into the *lgaldh* mutant (SALK_056664) background using reciprocal crosses. H) Diagnostic PCR of F2 generation from the cross indicated identified plants harboring l-GalDH-GFP in homozygous mutant background (arrowheads). Ascorbate content of this line is showed in Figure 2. E) Ascorbate content of the T3 generation of lines generated consisting of vtc2 mutants harboring VTC2p:VTC2-GFP. Three fully expanded leaves were collected on five-week-old T3 plants grown on soil in a growth chamber under short-day conditions (10 h light at 22°C/14 h dark at 19°C), at 60% humidity and 200 µmol m^−2^ s^−1^ light intensity. Different letters denote statistically significant differences for ascorbate content (One-Way ANOVA, α=0,05). G) Immunoblot (α-GFP) analysis of VTC4-GFP T2 generation showing the six lines expressing VTC4-GFP protein (background WT). After the identification of lines harboring single insertions (T2 Hyg^R^:Hyg^S^=3, Χ^2^ (α=0.05)), we selected line L5 to introduce the VTC4p:VTC4-GFP construct into the *vtc4* mutant (SALK_077222) background using reciprocal crosses. H) Diagnostic PCR of F2 generation from the cross indicated identified plants harboring VTC4-GFP in homozygous mutant background (arrowheads). Ascorbate content of this line is showed in Figure 2.

**Supplementary Figure 4.**
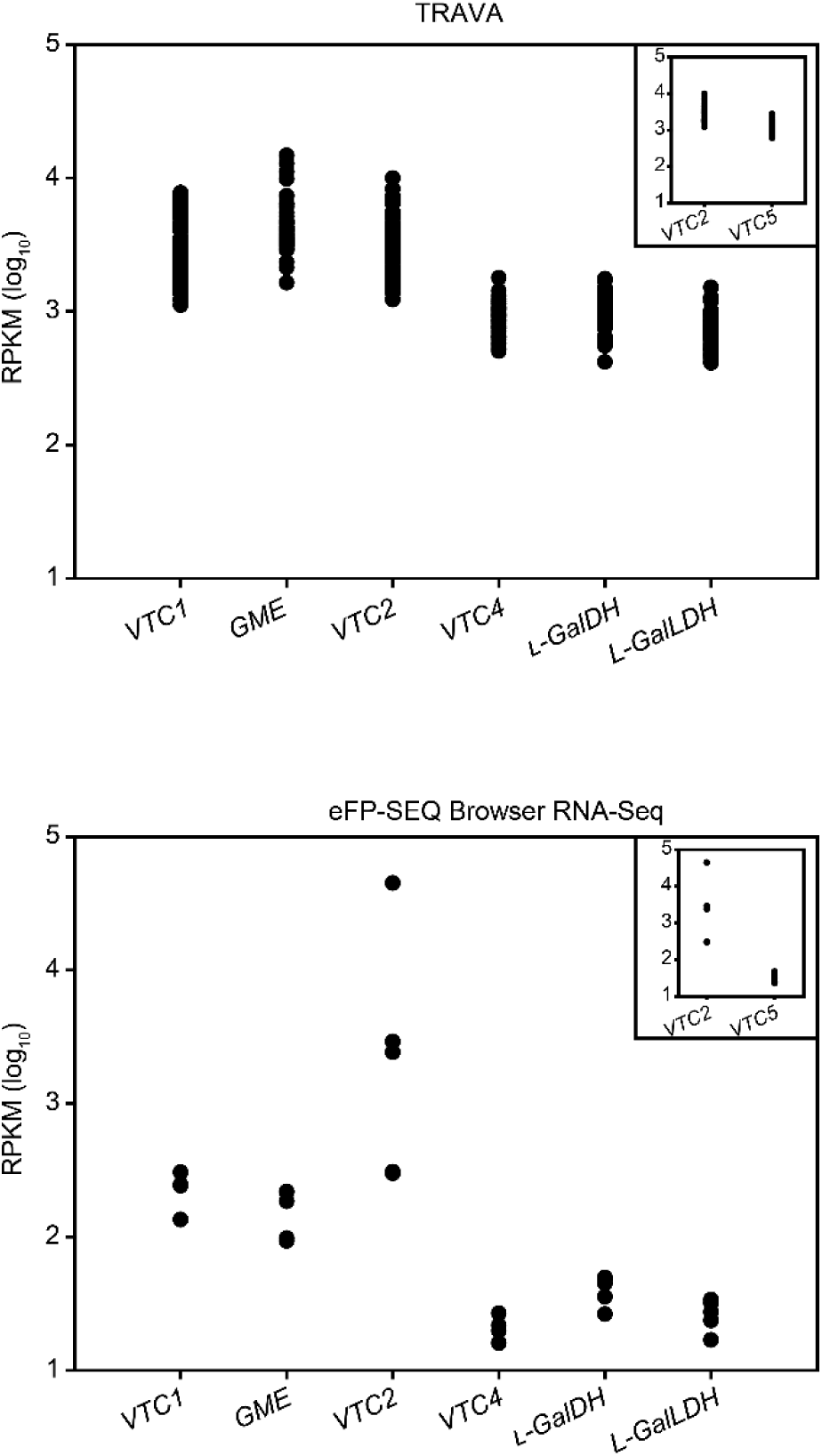
VTC2 mRNA levels are similar to those of VTC1 and GME, although VTC2 protein is hardly detectable. Data were taken from two different public RNA sequencing (RNA-seq) data sets: Transcriptome Variation Analysis (TRAVA, travadb.org; Klepikova et al., 2016) and the eFP-Seq Browser (https://bar.utoronto.ca/eFP-Seq_Browser/; Sullivan et al., 2019) corresponding to different adult plants’ vegetative and green tissues, respectively.

**Supplementary Figure 5.**
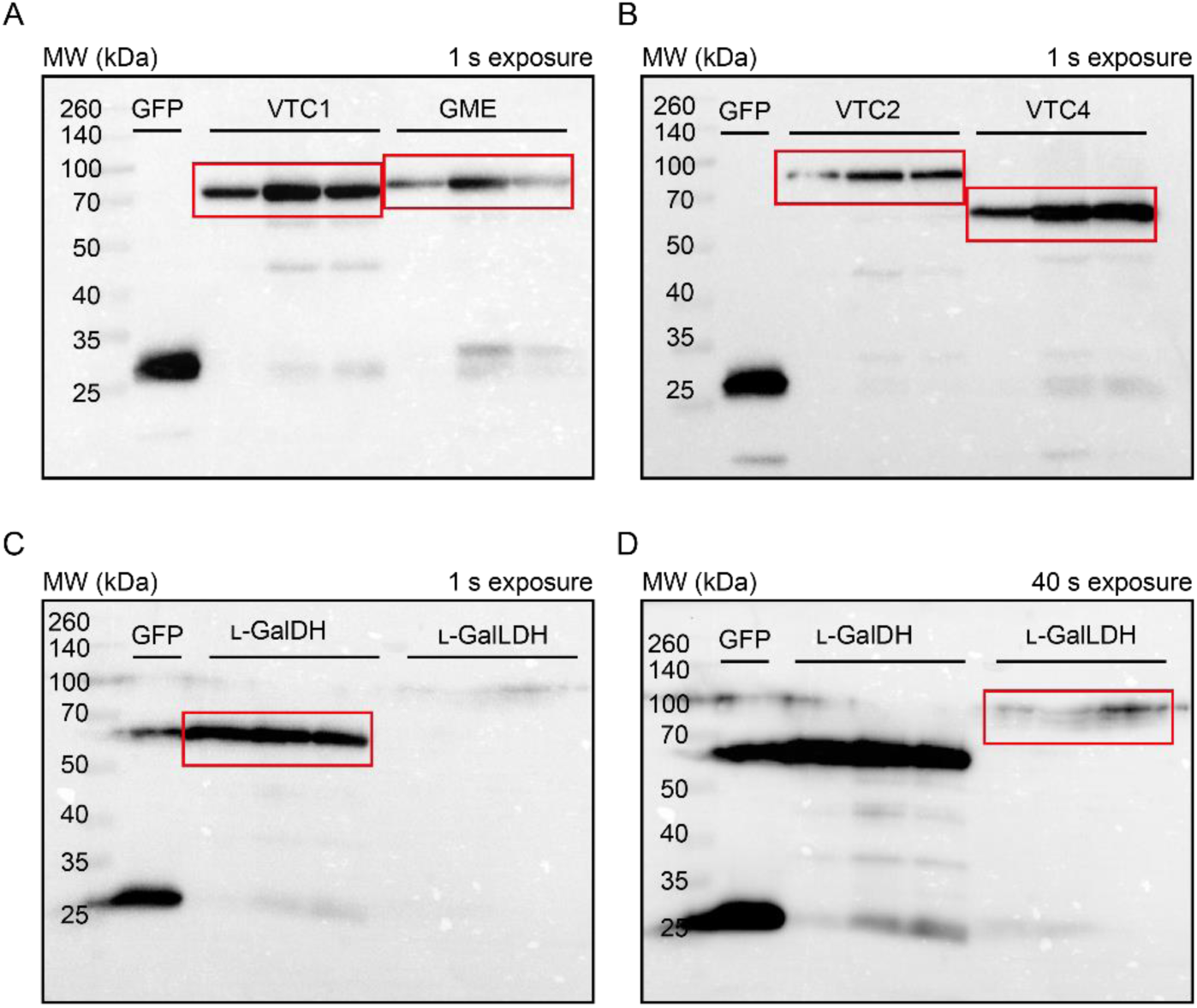
Complete blots corresponding to Figure 2B (red rectangle). Blots A, B and C have the same time of exposure using an ultra-sensitive enhanced chemiluminescent substrate for low femtogram protein level detection. Blot C and D are the same membrane with different exposure times. Protein loads were calculated using FIJI from a first immunoblot (not shown) and diluted so the expression of GFP-tagged proteins in the first timepoint is similar among constructs.

**Supplementary Figure 6.**
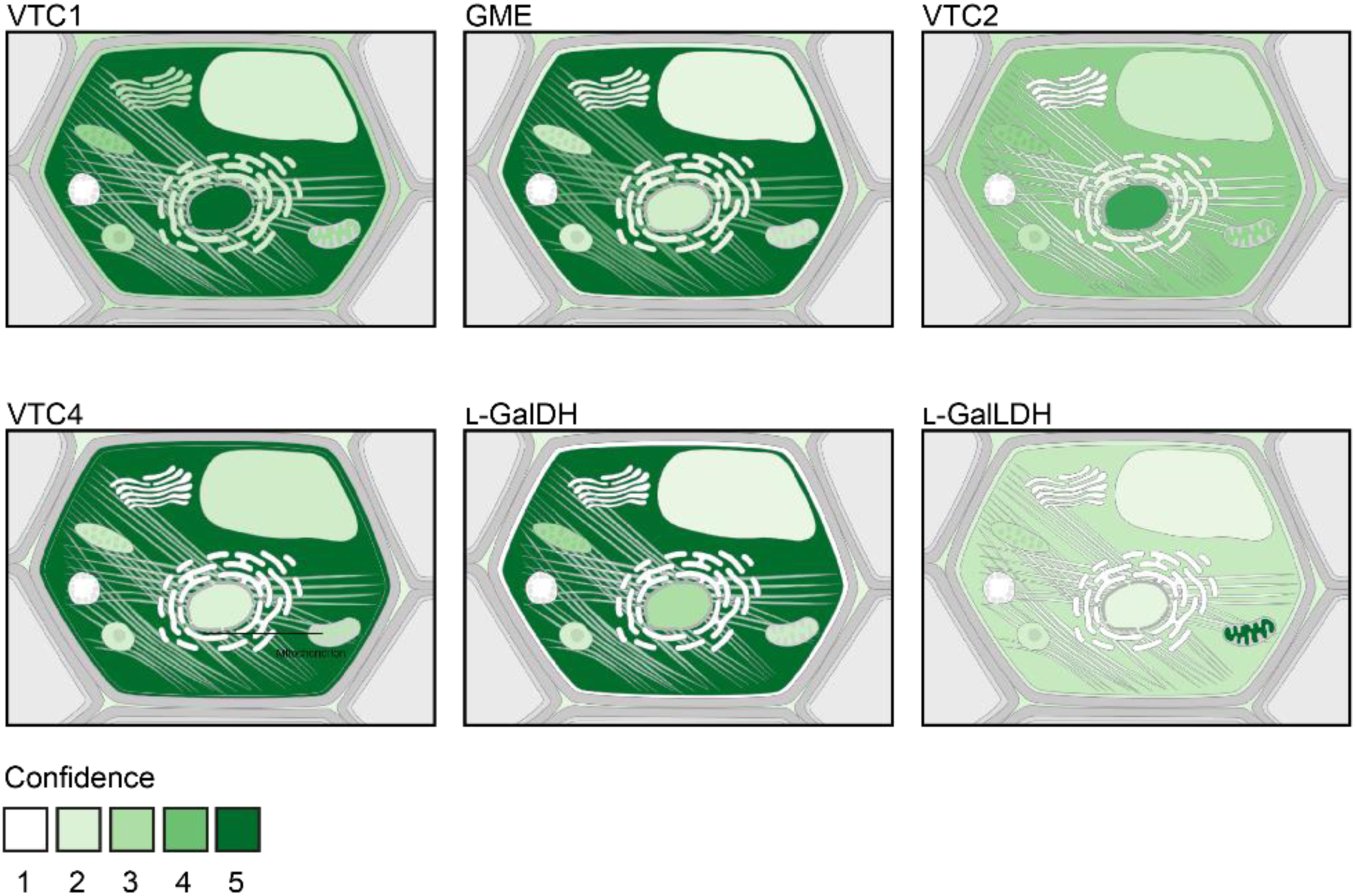
Predicted subcellular localization of ascorbate biosynthesis proteins in the plant cell according to COMPARTMENTS (https://compartments.jensenlab.org/Search; Binder et al., 2014).

**Supplementary Figure 7.**
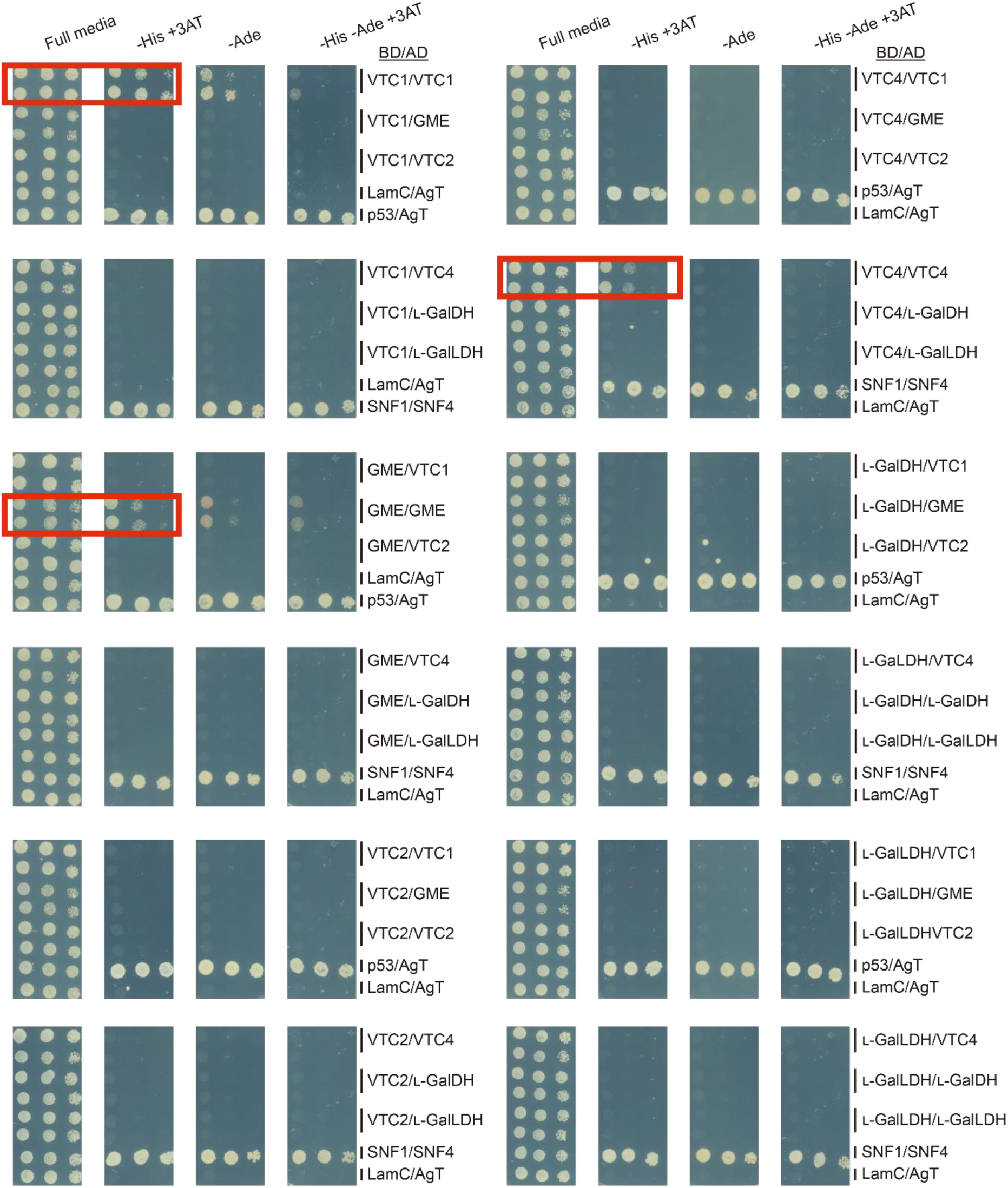
Yeast two-hybrid whole screening. Red rectangles correspond to the spotting presented in Figure 6A. As positive controls, we used SNF1/SNF4 (Fields and Song, 1989) and p53/SV40-Large AgT (AgT; Iwabuchi et al., 1993; Li and Fields, 1993) interactions. As a negative control, we included LamC/AgT cotransformation (Bartel et al., 1993; Ye and Worman, 1995). His: histidine, 3-AT: 3-amino-1,2,4-triazole, Ade: adenine.

**Supplementary Figure 8.**
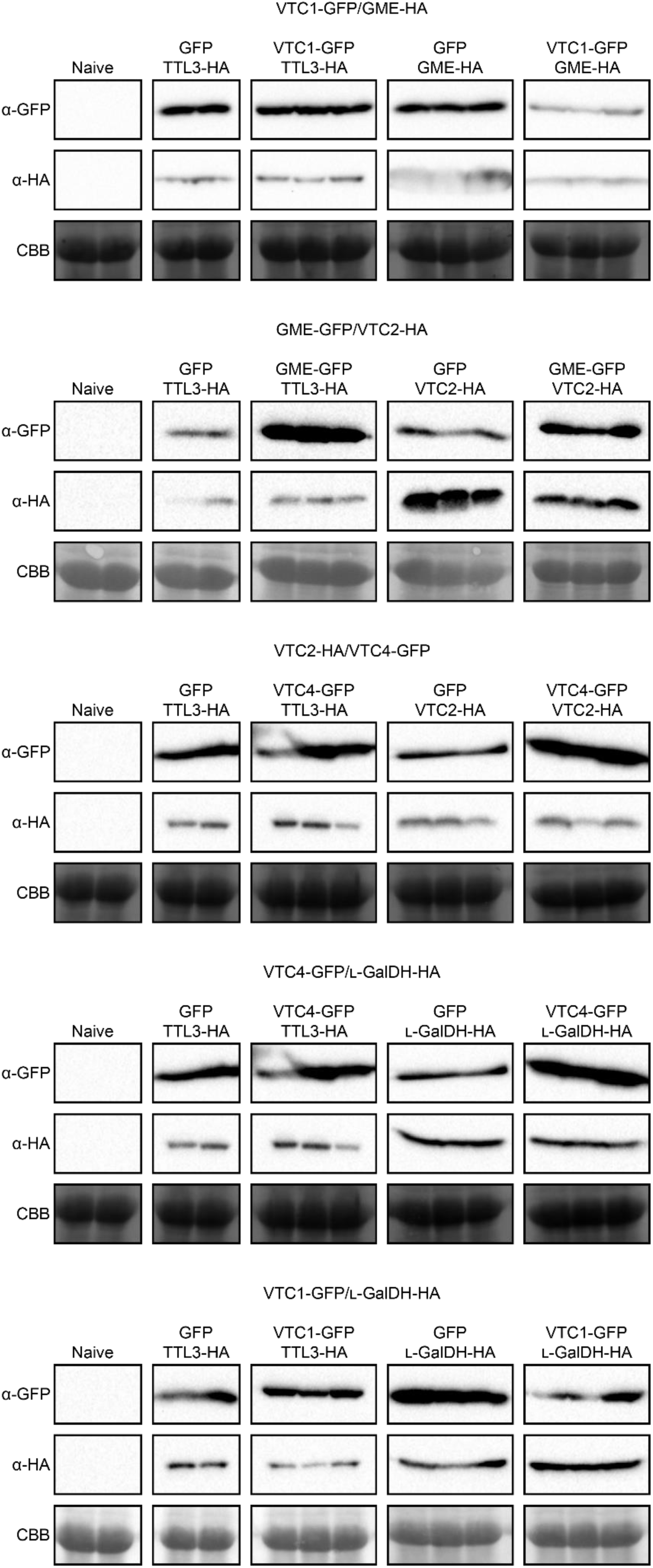
Immunoblot (α-GFP) of N. benthamiana leaves crude extracts following agroinfiltration of above-indicated combinations of proteins (also see Appendix 1). Two leaves of 2 or 3 independent plants were infiltrated, using GFP or TTL3-HA (an HA-tagged protein non-related to ascorbate synthesis) to rise optical density up to 1.

Supplementary Table 1. Infiltration of *Nicotiana benthamiana* corresponding to Figure 3C-H. * Data transformed with log10 x to accomplish normality and homoscedasticity required for ANOVA.

Supplementary Table 2. Prediction of transit peptides and nuclear localization signals within VTC genes. Values in the brackets are (probability | residues involved). Results obtained from LOCALIZER (http://localizer.csiro.au/; Sperschneider et al., 2017)

Supplementary Table 3. Initial conditions for the COPASI kinetic model of ascorbate biosynthesis and turnover.

Supplementary Table 4. Flux control coefficients for ascorbate biosynthesis and turnover with various strengths of non-competitive feedback inhibition at the GDP-l-Gal phosphorylase step (Ki GGP). Values were calculated in COPASI (Hoops et al., 2006) using conditions indicated in Supplementary Table 3.

Supplementary Table 5. Concentration control coefficients at each step for pathway intermediates with various strengths of non-competitive feedback inhibition at the GDP-l-Gal phosphorylase step (Ki GGP). Values were calculated in COPASI (Hoops et al., 2006) using conditions indicated in Supplementary Table 3.

Supplementary Table 6. Flux (A) and concentration (B) control coefficients at each step with competitive feedback inhibition of PMI and l-GalDH relaxed to 100 mM (compare Supplementary Tables 4 and 5 for published Ki values). Values were calculated in COPASI (Hoops et al., 2006) using conditions indicated in Supplementary Table 3.

Supplementary Table 7. The effect of varying relative enzyme activity (V or Vf) on the concentration of pathway intermediates with various strengths of non-competitive feedback inhibition at the GDP-l-Gal phosphorylase step (Ki GGP). Values were calculated in COPASI (Hoops et al., 2006) using conditions indicated in Supplementary Table 3.

Material and Methods Supplementary Table 1. Arabidopsis mutants and lines used in this work.

Material and Methods Supplementary Table 2. Arabidopsis stable lines used in this work.

Material and Methods Supplementary Table 3. A) Sizes of constructs generated in this work using the vectors listed in B) and the resultant construct.

## Acknowledgements

We acknowledge Glenda Gillaspy (Department of Biochemistry, Virginia Tech) for kindly providing us the *vtc4-4* mutant used in this work.

## Literature cited

Ali, B., Pantha, S., Acharya, R., Ueda, Y., Wu, L.B., Ashrafuzzaman, M., Ishizaki, T., Wissuwa, M., Bulley, S., Frei, M., 2019. Enhanced ascorbate level improves multi-stress tolerance in a widely grown indica rice variety without compromising its agronomic characteristics. J. Plant Physiol. https://doi.org/10.1016/j.jplph.2019.152998

Altschul, S.F., Madden, T.L., Schäffer, A.A., Zhang, J., Zhang, Z., Miller, W., Lipman, D.J., 1997. Gapped BLAST and PSI-BLAST: A new generation of protein database search programs. Nucleic Acids Res. https://doi.org/10.1093/nar/25.17.3389

Amorim-Silva, V., García-Moreno, Á., Castillo, A.G., Lakhssassi, N., Del Valle, A.E., Pérez-Sancho, J., Li, Y., Posé, D., Pérez-Rodriguez, J., Lin, J., Valpuesta, V., Borsani, O., Zipfel, C., Macho, A.P., Botella, M.A., 2019. TTL proteins scaffold brassinosteroid signaling components at the plasma membrane to optimize signal transduction in arabidopsis. Plant Cell. https://doi.org/10.1105/tpc.19.00150

Asada, K., 1999. THE WATER-WATER CYCLE IN CHLOROPLASTS: Scavenging of Active Oxygens and Dissipation of Excess Photons. Annu. Rev. Plant Physiol. Plant Mol. Biol. 50, 601–639. https://doi.org/10.1146/annurev.arplant.50.1.601

Badejo, A.A., Jeong, S.T., Goto-Yamamoto, N., Esaka, M., 2007. Cloning and expression of GDP-d-mannose pyrophosphorylase gene and ascorbic acid content of acerola (Malpighia glabra L.) fruit at ripening stages. Plant Physiol. Biochem. https://doi.org/10.1016/j.plaphy.2007.07.003

Badejo, A.A., Tanaka, N., Esaka, M., 2008. Analysis of GDP-D-Mannose Pyrophosphorylase Gene Promoter from Acerola (Malpighia glabra) and Increase in Ascorbate Content of Transgenic Tobacco Expressing the Acerola Gene. Plant Cell Physiol. 49, 126–132. https://doi.org/10.1093/pcp/pcm164

Bartel, P., Chien, C.T., Sternglanz, R., Fields, S., 1993. Elimination of false positives that arise in using the two-hybrid system. Biotechniques 14, 920–924.

Bartoli, C.G., 2006. Inter-relationships between light and respiration in the control of ascorbic acid synthesis and accumulation in Arabidopsis thaliana leaves. J. Exp. Bot. 57, 1621–1631. https://doi.org/10.1093/jxb/erl005

Binder, J.X., Pletscher-Frankild, S., Tsafou, K., Stolte, C., O’Donoghue, S.I., Schneider, R., Jensen, L.J., 2014. COMPARTMENTS: unification and visualization of protein subcellular localization evidence. Database 2014, bau012–bau012. https://doi.org/10.1093/database/bau012

Boukouris, A.E., Zervopoulos, S.D., Michelakis, E.D., 2016. Metabolic Enzymes Moonlighting in the Nucleus: Metabolic Regulation of Gene Transcription. Trends Biochem. Sci. 41, 712–730. https://doi.org/10.1016/j.tibs.2016.05.013

Bulley, S., Laing, W., 2016. The regulation of ascorbate biosynthesis. Curr. Opin. Plant Biol. 33, 15–22. https://doi.org/10.1016/j.pbi.2016.04.010

Bulley, S., Wright, M., Rommens, C., Yan, H., Rassam, M., Lin-Wang, K., Andre, C., Brewster, D., Karunairetnam, S., Allan, A.C., Laing, W.A., 2012. Enhancing ascorbate in fruits and tubers through over-expression of the L-galactose pathway gene GDP-L-galactose phosphorylase. Plant Biotechnol. J. 10, 390–397. https://doi.org/10.1111/j.1467-7652.2011.00668.x

Bulley, S.M., Rassam, M., Hoser, D., Otto, W., Schünemann, N., Wright, M., MacRae, E., Gleave, A., Laing, W., 2009. Gene expression studies in kiwifruit and gene over-expression in Arabidopsis indicates that GDP-L-galactose guanyltransferase is a major control point of vitamin C biosynthesis. J. Exp. Bot. 60, 765–778. https://doi.org/10.1093/jxb/ern327

Clough, S.J., Bent, A.F., 1998. Floral dip: a simplified method forAgrobacterium-mediated transformation ofArabidopsis thaliana. Plant J. 16, 735–743. https://doi.org/10.1046/j.1365-313x.1998.00343.x

Conklin, P.L., Gatzek, S., Wheeler, G.L., Dowdle, J., Raymond, M.J., Rolinski, S., Isupov, M., Littlechild, J.A., Smirnoff, N., 2006. Arabidopsis thaliana VTC4 encodes L-galactose-1-P phosphatase, a plant ascorbic acid biosynthetic enzyme. J. Biol. Chem. 281, 15662–15670. https://doi.org/10.1074/jbc.M601409200

Conklin, P.L., Norris, S.R., Wheeler, G.L., Williams, E.H., Smirnoff, N., Last, R.L., 1999. Genetic evidence for the role of GDP-mannose in plant ascorbic acid (vitamin C) biosynthesis. Proc. Natl. Acad. Sci. 96, 4198–4203. https://doi.org/10.1073/pnas.96.7.4198

Conklin, P.L., Pallanca, J.E., Last, R.L., Smirnoff, N., 1997. L-ascorbic acid metabolism in the ascorbate-deficient arabidopsis mutant vtc1. Plant Physiol. 115, 1277–85. https://doi.org/10.1016/s0168-9452(03)00277-2

Conklin, P.L., Saracco, S.A., Norris, S.R., Last, R.L., 2000. Identification of ascorbic acid-deficient Arabidopsis thaliana mutants. Genetics 154, 847–56.

Conklin, P.L., Willlams, E.H., Last, R.L., 1996. Environmental stress sensitivity of an ascorbic acid-deficient Arabidopsis mutant. Proc. Natl. Acad. Sci. 93, 9970–9974.

Cronje, C., George, G.M., Fernie, A.R., Bekker, J., Kossmann, J., Bauer, R., 2012. Manipulation of L-ascorbic acid biosynthesis pathways in Solanum lycopersicum: elevated GDP-mannose pyrophosphorylase activity enhances L-ascorbate levels in red fruit. Planta 235, 553–564. https://doi.org/10.1007/s00425-011-1525-6

Davey, M.W., Kenis, K., Keulemans, J., 2006. Genetic control of fruit vitamin C contents. Plant Physiol. https://doi.org/10.1104/pp.106.083279

DeBolt, S., Cook, D.R., Ford, C.M., 2006. L-Tartaric acid synthesis from vitamin C in higher plants. Proc. Natl. Acad. Sci. 103, 5608–5613. https://doi.org/10.1073/pnas.0510864103

Debolt, S., Melino, V., Ford, C.M., 2007. Ascorbate as a Biosynthetic Precursor in Plants. Ann. Bot. 99, 3–8. https://doi.org/10.1093/aob/mcl236

Dewhirst, R.A., Clarkson, G.J.J., Rothwell, S.D., Fry, S.C., 2017. Novel insights into ascorbate retention and degradation during the washing and post-harvest storage of spinach and other salad leaves. Food Chem. https://doi.org/10.1016/j.foodchem.2017.04.082

Dewhirst, R.A., Fry, S.C., 2018. The oxidation of dehydroascorbic acid and 2,3-diketogulonate by distinct reactive oxygen species. Biochem. J. https://doi.org/10.1042/BCJ20180688

Dewhirst, R.A., Murray, L., Mackay, C.L., Sadler, I.H., Fry, S.C., 2020. Characterisation of the non-oxidative degradation pathway of dehydroascorbic acid in slightly acidic aqueous solution. Arch. Biochem. Biophys. 681, 108240. https://doi.org/10.1016/j.abb.2019.108240

Dowdle, J., Ishikawa, T., Gatzek, S., Rolinski, S., Smirnoff, N., 2007. Two genes in Arabidopsis thaliana encoding GDP-L-galactose phosphorylase are required for ascorbate biosynthesis and seedling viability. Plant J. https://doi.org/10.1111/j.1365-313X.2007.03266.x

Dunkley, T.P.J., Hester, S., Shadforth, I.P., Runions, J., Weimar, T., Hanton, S.L., Griffin, J.L., Bessant, C., Brandizzi, F., Hawes, C., Watson, R.B., Dupree, P., Lilley, K.S., 2006. Mapping the Arabidopsis organelle proteome. Proc. Natl. Acad. Sci. 103, 6518–6523. https://doi.org/10.1073/pnas.0506958103

Fell, D.A., 1992. Metabolic control analysis: a survey of its theoretical and experimental development. Biochem. J. 286, 313–330. https://doi.org/10.1042/bj2860313

Fenech, M., Amaya, I., Valpuesta, V., Botella, M.A., 2019. Vitamin C content in fruits: Biosynthesis and regulation. Front. Plant Sci. 9, 1–21. https://doi.org/10.3389/fpls.2018.02006

Fields, S., Song, O., 1989. A novel genetic system to detect protein–protein interactions. Nature 340, 245–246. https://doi.org/10.1038/340245a0

Franceschi, V.R., Tarlyn, N.M., 2002. L-Ascorbic Acid Is Accumulated in Source Leaf Phloem and Transported to Sink Tissues in Plants. Plant Physiol. 130, 649–656. https://doi.org/10.1104/pp.007062

Gao, Y., Badejo, A.A., Shibata, H., Sawa, Y., Maruta, T., Shigeoka, S., Page, M., Smirnoff, N., Ishikawa, T., 2011. Expression Analysis of the VTC2 and VTC5 Genes Encoding GDP-L-Galactose Phosphorylase, an Enzyme Involved in Ascorbate Biosynthesis, in Arabidopsis thaliana. Biosci. Biotechnol. Biochem. 75, 1783–1788. https://doi.org/10.1271/bbb.110320

Gatzek, S., Wheeler, G.L., Smirnoff, N., 2002. Antisense suppression of L-galactose dehydrogenase in Arabidopsis thaliana provides evidence for its role in ascorbate synthesis and reveals light modulated L-galactose synthesis. Plant J. https://doi.org/10.1046/j.1365-313X.2002.01315.x

Gietz, R.D., Schiestl, R.H., 1995. Transforming yeast with DNA. Methods Mol. Cell. Biol. 5, 255–269.

Gilbert, L., Alhagdow, M., Nunes-Nesi, A., Quemener, B., Guillon, F., Bouchet, B., Faurobert, M., Gouble, B., Page, D., Garcia, V., Petit, J., Stevens, R., Causse, M., Fernie, A.R., Lahaye, M., Rothan, C., Baldet, P., 2009. GDP-d-mannose 3,5-epimerase (GME) plays a key role at the intersection of ascorbate and non-cellulosic cell-wall biosynthesis in tomato. Plant J. 60, 499–508. https://doi.org/10.1111/j.1365-313X.2009.03972.x

Green, M.A., Fry, S.C., 2005. Vitamin C degradation in plant cells via enzymatic hydrolysis of 4-O-oxalyl-l-threonate. Nature 433, 83–87. https://doi.org/10.1038/nature03172

Harapanhalli, R.S., Howell, R.W., Rao, D. V., 1993. Testicular and plasma ascorbic acid levels in mice following dietary intake: a high-performance liquid chromatographic analysis. J. Chromatogr. B Biomed. Sci. Appl. https://doi.org/10.1016/0378-4347(93)80314-T

Heazlewood, J.L., Tonti-Filippini, J.S., Gout, A.M., Day, D.A., Whelan, J., Millar, A.H., 2004. Experimental Analysis of the Arabidopsis Mitochondrial Proteome Highlights Signaling and Regulatory Components, Provides Assessment of Targeting Prediction Programs, and Indicates Plant-Specific Mitochondrial Proteins. Plant Cell 16, 241–256. https://doi.org/10.1105/tpc.016055

Hoeberichts, F.A., Vaeck, E., Kiddle, G., Coppens, E., Van De Cotte, B., Adamantidis, A., Ormenese, S., Foyer, C.H., Zabeau, M., Inzé, D., Périlleux, C., Van Breusegem, F., Vuylsteke, M., 2008. A temperature-sensitive mutation in the Arabidopsis thaliana phosphomannomutase gene disrupts protein glycosylation and triggers cell death. J. Biol. Chem. 283, 5708–5718. https://doi.org/10.1074/jbc.M704991200

Hoops, S., Gauges, R., Lee, C., Pahle, J., Simus, N., Singhal, M., Xu, L., Mendes, P., Kummer, U., 2006. COPASI - A COmplex PAthway SImulator. Bioinformatics. https://doi.org/10.1093/bioinformatics/btl485

Iwabuchi, K., Li, B., Bartel, P., Fields, S., 1993. Use of the two-hybrid system to identify the domain of p53 involved in oligomerization. Oncogene 8, 1693–6.

Klepikova, A. V., Kasianov, A.S., Gerasimov, E.S., Logacheva, M.D., Penin, A.A., 2016. A high resolution map of the Arabidopsis thaliana developmental transcriptome based on RNA-seq profiling. Plant J. 88, 1058–1070. https://doi.org/10.1111/tpj.13312

Laing, W., Norling, C., Brewster, D., Wright, M., Bulley, S., 2017. Ascorbate concentration in *Arabidopsis thaliana* and expression of ascorbate related genes using RNAseq in response to light and the diurnal cycle. bioRxiv 138008. https://doi.org/10.1101/138008

Laing, W. A., Bulley, S., Wright, M., Cooney, J., Jensen, D., Barraclough, D., MacRae, E., 2004. A highly specific L-galactose-1-phosphate phosphatase on the path to ascorbate biosynthesis. Proc. Natl. Acad. Sci. 101, 16976–16981. https://doi.org/10.1073/pnas.0407453101

Laing, William A, Frearson, N., Bulley, S., MacRae, E., 2004. Kiwifruit L-galactose dehydrogenase: molecular, biochemical and physiological aspects of the enzyme. Funct. Plant Biol. 31, 1015–1025. https://doi.org/10.1071/FP04090

Laing, W.A., Martínez-Sánchez, M., Wright, M.A., Bulley, S.M., Brewster, D., Dare, A.P., Rassam, M., Wang, D., Storey, R., Macknight, R.C., Hellens, R.P., 2015. An Upstream Open Reading Frame Is Essential for Feedback Regulation of Ascorbate Biosynthesis in Arabidopsis. Plant Cell. https://doi.org/10.1105/tpc.114.133777

Laing, W.A., Wright, M.A., Cooney, J., Bulley, S.M., 2007. The missing step of the L-galactose pathway of ascorbate biosynthesis in plants, an L-galactose guanyltransferase, increases leaf ascorbate content. Proc. Natl. Acad. Sci. 104, 9534–9539.

Li, B., Fields, S., 1993. Identification of mutations in p53 that affect its binding to SV40 large T antigen by using the yeast two-hybrid system. FASEB J. https://doi.org/10.1096/fasebj.7.10.8344494

Li, J., Liang, D., Li, M., Ma, F., 2013. Light and abiotic stresses regulate the expression of GDP-L-galactose phosphorylase and levels of ascorbic acid in two kiwifruit genotypes via light-responsive and stress-inducible cis-elements in their promoters. Planta 238, 535–547. https://doi.org/10.1007/s00425-013-1915-z

Li, S., Wang, J., Yu, Y., Wang, F., Dong, J., Huang, R., 2016. D27E mutation of VTC1 impairs the interaction with CSN5B and enhances ascorbic acid biosynthesis and seedling growth in Arabidopsis. Plant Mol. Biol. 92, 473–482. https://doi.org/10.1007/s11103-016-0525-0

Li, T., Yang, X., Yu, Y., Si, X., Zhai, X., Zhang, H., Dong, W., Gao, C., Xu, C., 2018. Domestication of wild tomato is accelerated by genome editing. Nat. Biotechnol. https://doi.org/10.1038/nbt.4273

Li, X., Ye, J., Munir, S., Yang, T., Chen, W., Liu, G., Zheng, W., Zhang, Y., 2019. Biosynthetic Gene Pyramiding Leads to Ascorbate Accumulation with Enhanced Oxidative Stress Tolerance in Tomato. Int. J. Mol. Sci. 20, 1558. https://doi.org/10.3390/ijms20071558

Lim, B., Smirnoff, N., Cobbett, C.S., Golz, J.F., 2016. Ascorbate-Deficient vtc2 Mutants in Arabidopsis Do Not Exhibit Decreased Growth. Front. Plant Sci. 7, 1–9. https://doi.org/10.3389/fpls.2016.01025

Linster, C.L., Gomez, T.A., Christensen, K.C., Adler, L.N., Young, B.D., Brenner, C., Clarke, S.G., 2007. Arabidopsis VTC2 encodes a GDP-L-galactose phosphorylase, the last unknown enzyme in the smirnoff-wheeler pathway to ascorbic acid in plants. J. Biol. Chem. https://doi.org/10.1074/jbc.M702094200

Lorence, A., Chevone, B.I., Mendes, P., Nessler, C.L., 2004. myo-Inositol Oxygenase Offers a Possible Entry Point into Plant Ascorbate Biosynthesis. Plant Physiol. 134, 1200–1205. https://doi.org/10.1104/pp.103.033936

Lukowitz, W., Nickle, T.C., Meinke, D.W., Last, R.L., Conklin, P.L., Somerville, C.R., 2001. Arabidopsis cyt1 mutants are deficient in a mannose-1-phosphate guanylyltransferase and point to a requirement of N-linked glycosylation for cellulose biosynthesis. Proc. Natl. Acad. Sci. 98, 2262–2267. https://doi.org/10.1073/pnas.051625798

Marchler-Bauer, A., Bo, Y., Han, L., He, J., Lanczycki, C.J., Lu, S., Chitsaz, F., Derbyshire, M.K., Geer, R.C., Gonzales, N.R., Gwadz, M., Hurwitz, D.I., Lu, F., Marchler, G.H., Song, J.S., Thanki, N., Wang, Z., Yamashita, R.A., Zhang, D., Zheng, C., Geer, L.Y., Bryant, S.H., 2017. CDD/SPARCLE: functional classification of proteins via subfamily domain architectures. Nucleic Acids Res. 45, D200–D203. https://doi.org/10.1093/nar/gkw1129

Marchler-Bauer, A., Bryant, S.H., 2004. CD-Search: protein domain annotations on the fly. Nucleic Acids Res. 32, W327–W331. https://doi.org/10.1093/nar/gkh454

Marchler-Bauer, A., Derbyshire, M.K., Gonzales, N.R., Lu, S., Chitsaz, F., Geer, L.Y., Geer, R.C., He, J., Gwadz, M., Hurwitz, D.I., Lanczycki, C.J., Lu, F., Marchler, G.H., Song, J.S., Thanki, N., Wang, Z., Yamashita, R.A., Zhang, D., Zheng, C., Bryant, S.H., 2015. CDD: NCBI’s conserved domain database. Nucleic Acids Res. https://doi.org/10.1093/nar/gku1221

Marchler-Bauer, A., Lu, S., Anderson, J.B., Chitsaz, F., Derbyshire, M.K., DeWeese-Scott, C., Fong, J.H., Geer, L.Y., Geer, R.C., Gonzales, N.R., Gwadz, M., Hurwitz, D.I., Jackson, J.D., Ke, Z., Lanczycki, C.J., Lu, F., Marchler, G.H., Mullokandov, M., Omelchenko, M. V., Robertson, C.L., Song, J.S., Thanki, N., Yamashita, R.A., Zhang, D., Zhang, N., Zheng, C., Bryant, S.H., 2011. CDD: a Conserved Domain Database for the functional annotation of proteins. Nucleic Acids Res. 39, D225–D229. https://doi.org/10.1093/nar/gkq1189

Maruta, T., Yonemitsu, M., Yabuta, Y., Tamoi, M., Ishikawa, T., Shigeoka, S., 2008. Arabidopsis phosphomannose isomerase 1, but not phosphomannose isomerase 2, is essential for ascorbic acid biosynthesis. J. Biol. Chem. 283, 28842–28851. https://doi.org/10.1074/jbc.M805538200

Mellidou, I., Chagne, D., Laing, W.A., Keulemans, J., Davey, M.W., 2012. Allelic Variation in Paralogs of GDP-l-Galactose Phosphorylase Is a Major Determinant of Vitamin C Concentrations in Apple Fruit. Plant Physiol. 160, 1613–1629. https://doi.org/10.1104/pp.112.203786

Mieda, T., Yabuta, Y., Rapolu, M., Motoki, T., Takeda, T., Yoshimura, K., Ishikawa, T., Shigeoka, S., 2004. Feedback inhibition of spinach L-galactose dehydrogenase by L-ascorbate. Plant Cell Physiol. 45, 1271–1279. https://doi.org/10.1093/pcp/pch152

Müller-Moulé, P., 2008. An expression analysis of the ascorbate biosynthesis enzyme VTC2. Plant Mol. Biol. 68, 31–41. https://doi.org/10.1007/s11103-008-9350-4

Nakagawa, T., Kurose, T., Hino, T., Tanaka, K., Kawamukai, M., Niwa, Y., Toyooka, K., Matsuoka, K., Jinbo, T., Kimura, T., 2007. Development of series of gateway binary vectors, pGWBs, for realizing efficient construction of fusion genes for plant transformation. J. Biosci. Bioeng. https://doi.org/10.1263/jbb.104.34

Nourbakhsh, A., Collakova, E., Gillaspy, G.E., 2015. Characterization of the inositol monophosphatase gene family in Arabidopsis. Front. Plant Sci. 5, 1–14. https://doi.org/10.3389/fpls.2014.00725

Østergaard, J., Persiau, G., Davey, M.W., Bauw, G., Van Montagu, M., 1997. Isolation of a cDNA coding for L-galactono-gamma-lactone dehydrogenase, an enzyme involved in the biosynthesis of ascorbic acid in plants. Purification, characterization, cDNA cloning, and expression in yeast. J. Biol. Chem. 272, 30009–16.

Page, M., Sultana, N., Paszkiewicz, K., Florance, H., Smirnoff, N., 2012. The influence of ascorbate on anthocyanin accumulation during high light acclimation in Arabidopsis thaliana: further evidence for redox control of anthocyanin synthesis. Plant. Cell Environ. 35, 388–404. https://doi.org/10.1111/j.1365-3040.2011.02369.x

Pallanca, J.E., Smirnoff, N., 2000. The control of ascorbic acid synthesis and turnover in pea seedlings. J. Exp. Bot. 51, 669–674. https://doi.org/10.1093/jexbot/51.345.669

Pineau, B., Layoune, O., Danon, A., De Paepe, R., 2008. L-galactono-1,4-lactone dehydrogenase is required for the accumulation of plant respiratory complex I. J. Biol. Chem. 283, 32500–32505. https://doi.org/10.1074/jbc.M805320200

Plumb, W., Townsend, A.J., Rasool, B., Alomrani, S., Razak, N., Karpinska, B., Ruban, A. V., Foyer, C.H., 2018. Ascorbate-mediated regulation of growth, photoprotection, and photoinhibition in Arabidopsis thaliana. J. Exp. Bot. https://doi.org/10.1093/jxb/ery170

Qi, T., Liu, Z., Fan, M., Chen, Y., Tian, H., Wu, D., Gao, H., Ren, C., Song, S., Xie, D., 2017. GDP-D-mannose epimerase regulates male gametophyte development, plant growth and leaf senescence in Arabidopsis. Sci. Rep. 7, 10309. https://doi.org/10.1038/s41598-017-10765-5

Reiter, W.D., Vanzin, G.F., 2001. Molecular genetics of nucleotide sugar interconversion pathways in plants. Plant Mol. Biol. 47, 95–113.

Sawake, S., Tajima, N., Mortimer, J.C., Lao, J., Ishikawa, T., Yu, X., Yamanashi, Y., Yoshimi, Y., Kawai-Yamada, M., Dupree, P., Tsumuraya, Y., Kotake, T., 2015. KONJAC1 and 2 Are Key Factors for GDP-Mannose Generation and Affect l-Ascorbic Acid and Glucomannan Biosynthesis in Arabidopsis. Plant Cell. https://doi.org/10.1105/tpc.15.00379

Schimmeyer, J., Bock, R., Meyer, E.H., 2016. L-Galactono-1,4-lactone dehydrogenase is an assembly factor of the membrane arm of mitochondrial complex I in Arabidopsis. Plant Mol. Biol. 90, 117–126. https://doi.org/10.1007/s11103-015-0400-4

Schindelin, J., Arganda-Carreras, I., Frise, E., Kaynig, V., Longair, M., Pietzsch, T., Preibisch, S., Rueden, C., Saalfeld, S., Schmid, B., Tinevez, J.-Y., White, D.J., Hartenstein, V., Eliceiri, K., Tomancak, P., Cardona, A., 2012. Fiji: an open-source platform for biological-image analysis. Nat. Methods 9, 676–682. https://doi.org/10.1038/nmeth.2019

Schneider, C.A., Rasband, W.S., Eliceiri, K.W., 2012. NIH Image to ImageJ: 25 years of image analysis. Nat. Methods 9, 671–675. https://doi.org/10.1038/nmeth.2089

Smirnoff, N., 2019. Engineering of metabolic pathways using synthetic enzyme complexes. Plant Physiol. https://doi.org/10.1104/pp.18.01280

Smirnoff, N., 2018. Ascorbic acid metabolism and functions: A comparison of plants and mammals. Free Radic. Biol. Med. 0–1. https://doi.org/10.1016/j.freeradbiomed.2018.03.033

Sperschneider, J., Catanzariti, A.M., Deboer, K., Petre, B., Gardiner, D.M., Singh, K.B., Dodds, P.N., Taylor, J.M., 2017. LOCALIZER: Subcellular localization prediction of both plant and effector proteins in the plant cell. Sci. Rep. https://doi.org/10.1038/srep44598

Sullivan, A., Purohit, P.K., Freese, N.H., Pasha, A., Esteban, E., Waese, J., Wu, A., Chen, M., Chin, C.Y., Song, R., Watharkar, S.R., Chan, A.P., Krishnakumar, V., Vaughn, M.W., Town, C., Loraine, A.E., Provart, N.J., 2019. An ‘<scp>eFP</scp> -Seq Browser’ for visualizing and exploring <scp>RNA</scp> sequencing data. Plant J. 100, 641–654. https://doi.org/10.1111/tpj.14468

Sweetlove, L.J., Fernie, A.R., 2018. The role of dynamic enzyme assemblies and substrate channelling in metabolic regulation. Nat. Commun. 9, 2136. https://doi.org/10.1038/s41467-018-04543-8

Terai, Y., Ueno, H., Ogawa, T., Sawa, Y., Miyagi, A., Kawai-Yamada, M., Ishikawa, T., Maruta, T., 2020. Dehydroascorbate Reductases and Glutathione Set a Threshold for High-Light-Induced Ascorbate Accumulation. Plant Physiol. https://doi.org/10.1104/pp.19.01556

Torabinejad, J., Donahue, J.L., Gunesekera, B.N., Allen-Daniels, M.J., Gillaspy, G.E., 2009. VTC4 Is a Bifunctional Enzyme That Affects Myoinositol and Ascorbate Biosynthesis in Plants. Plant Physiol. 150, 951–961. https://doi.org/10.1104/pp.108.135129

Truffault, V., Fry, S.C., Stevens, R.G., Edinburgh, T., Wall, C., Building, D.R., Gautier, H., Edinburgh, T., Wall, C., Building, D.R., 2017. Ascorbate degradation in tomato leads to accumulation of oxalate, threonate and oxalyl threonate. Plant J. 89, 996–1008. https://doi.org/10.1111/tpj.13439

Urzica, E.I., Adler, L.N., Page, M.D., Linster, C.L., Arbing, M.A., Casero, D., Pellegrini, M., Merchant, S.S., Clarke, S.G., 2012. Impact of oxidative stress on ascorbate biosynthesis in Chlamydomonas via regulation of the VTC2 gene encoding a GDP-L-galactose phosphorylase. J. Biol. Chem. 287, 14234–14245. https://doi.org/10.1074/jbc.M112.341982

Voxeur, A., Gilbert, L., Rihouey, C., Driouich, A., Rothan, C., Baldet, P., Lerouge, P., 2011. Silencing of the GDP-D-mannose 3,5-epimerase affects the structure and cross-linking of the pectic polysaccharide rhamnogalacturonan II and plant growth in tomato. J. Biol. Chem. 286, 8014–8020. https://doi.org/10.1074/jbc.M110.198614

Wang, J., Yu, Y., Zhang, Z., Quan, R., Zhang, H., Ma, L., Deng, X.W., Huang, R., 2013. Arabidopsis CSN5B Interacts with VTC1 and Modulates Ascorbic Acid Synthesis. Plant Cell. https://doi.org/10.1105/tpc.112.106880

Wang, H. Sen, Yu, C., Zhu, Z.J., Yu, X.C., 2011. Overexpression in tobacco of a tomato GMPase gene improves tolerance to both low and high temperature stress by enhancing antioxidation capacity. Plant Cell Rep. 30, 1029–1040. https://doi.org/10.1007/s00299-011-1009-y

Waszczak, C., Carmody, M., Kangasjärvi, J., 2018. Reactive Oxygen Species in Plant Signaling. Annu. Rev. Plant Biol. 69, 209–236. https://doi.org/10.1007/978-3-642-00390-5

Wheeler, G., Ishikawa, T., Pornsaksit, V., Smirnoff, N., 2015. Evolution of alternative biosynthetic pathways for vitamin C following plastid acquisition in photosynthetic eukaryotes. Elife 2015, 1–14. https://doi.org/10.7554/eLife.06369

Wheeler, G.L., Jones, M.A., Smirnoff, N., 1998. The biosynthetic pathway of vitamin C in higher plants. Nature 393, 365–369. https://doi.org/10.1038/30728

Wolucka, B.A., Van Montagu, M., 2003. GDP-Mannose 3′,5′-Epimerase Forms GDP-L-gulose, a Putative Intermediate for the de Novo Biosynthesis of Vitamin C in Plants. J. Biol. Chem. 278, 47483–47490. https://doi.org/10.1074/jbc.M309135200

Yabuta, Y., Mieda, T., Rapolu, M., Nakamura, A., Motoki, T., Maruta, T., Yoshimura, K., Ishikawa, T., Shigeoka, S., 2007. Light regulation of ascorbate biosynthesis is dependent on the photosynthetic electron transport chain but independent of sugars in Arabidopsis. J. Exp. Bot. 58, 2661–2671. https://doi.org/10.1093/jxb/erm124

Ye, Q., Worman, H.J., 1995. Protein-protein interactions between human nuclear lamins expressed in yeast. Exp. Cell Res. https://doi.org/10.1006/excr.1995.1230

Yoshimura, K., Nakane, T., Kume, S., Shiomi, Y., Maruta, T., Ishikawa, T., Shigeoka, S., 2014. Transient expression analysis revealed the importance of VTC2 expression level in light/dark regulation of ascorbate biosynthesis in Arabidopsis 78, 60–6. https://doi.org/10.1080/09168451.2014.877831

Zhang, C., Liu, J., Zhang, Y., Cai, X., Gong, P., Zhang, J., Wang, T., Li, H., Ye, Z., 2011. Overexpression of SlGMEs leads to ascorbate accumulation with enhanced oxidative stress, cold, and salt tolerance in tomato. Plant Cell Rep. 30, 389–398. https://doi.org/10.1007/s00299-010-0939-0

Zhang, G.Y., Liu, R.R., Zhang, C.Q., Tang, K.X., Sun, M.F., Yan, G.H., Liu, Q.Q., 2015. Manipulation of the rice L-galactose pathway: Evaluation of the effects of transgene overexpression on ascorbate accumulation and abiotic stress tolerance. PLoS One. https://doi.org/10.1371/journal.pone.0125870

Zhang, H., Si, X., Ji, X., Fan, R., Liu, J., Chen, K., Wang, D., Gao, C., 2018. Genome editing of upstream open reading frames enables translational control in plants. Nat. Biotechnol. 36, 894–898. https://doi.org/10.1038/nbt.4202

Zhang, Y., Beard, K.F.M., Swart, C., Bergmann, S., Krahnert, I., Nikoloski, Z., Graf, A., George Ratcliffe, R., Sweetlove, L.J., Fernie, A.R., Obata, T., 2017. Protein-protein interactions and metabolite channelling in the plant tricarboxylic acid cycle. Nat. Commun. 8. https://doi.org/10.1038/ncomms15212

Zhang, Y., Butelli, E., Martin, C., 2014. Engineering anthocyanin biosynthesis in plants. Curr. Opin. Plant Biol. https://doi.org/10.1016/j.pbi.2014.05.011

Zhou, Y., Tao, Q.C., Wang, Z.N., Fan, R., Li, Y., Sun, X.F., Tang, K.X., 2012. Engineering ascorbic acid biosynthetic pathway in Arabidopsis leaves by single and double gene transformation. Biol. Plant. 56, 451–457.

